# Hypoxia-inducible factor-2 (HIF2) regulates alveolar regeneration after repetitive injury

**DOI:** 10.1101/2023.09.17.557477

**Authors:** A. Scott McCall, Harikrishna Tanjore, Ankita Burman, Taylor Sherrill, Micah Chapman, Carla L. Calvi, Jane Camarata, Raphael P. Hunt, David Nichols, Nicholas E. Banovich, William E. Lawson, Jason J. Gokey, Jonathan A. Kropski, Timothy S. Blackwell

## Abstract

Idiopathic Pulmonary Fibrosis (IPF) is a progressive and often fatal chronic respiratory disease thought to result from repetitive injury and failed repair of the lung alveoli, and recent studies have identified a number of disease-emergent intermediate/transitional cell states in the IPF lung supporting this concept. In this study, we found that persistent activation of hypoxia-inducible factor (HIF)-signaling in airway-derived, repair-associated cell types/states is a hallmark of dysfunctional epithelial repair in the IPF lung epithelium and experimental models of recurrent lung epithelial injury. Disrupting Hif-signaling attenuated experimental lung fibrosis, reduced mucous-secretory cell polarization, and promoted functional alveolar regeneration following repetitive injury. Mouse and human organoid studies demonstrated that small-molecule-based HIF2 inhibition promoted alveolar epithelial cell proliferation and maturation while preventing the emergence of maladaptive intermediate/transitional states analogous to those in IPF. Together, these studies indicate that targeted HIF2-inhibition represents a novel and effective therapeutic strategy to promote functional lung regeneration, and could be readily translated into human studies of IPF and other chronic interstitial lung diseases with disease modifying effect.

**One sentence summary:** Inhibiting hypoxia-inducible-factor 2 (HIF2) promotes functional lung alveolar epithelial repair following recurrent injury.

## Introduction

Despite the development and widespread utilization of first-generation antifibrotic therapies(*1, 2*), the morbidity and mortality of Idiopathic Pulmonary Fibrosis (IPF) remains high(*3*) and there is substantial unmet need for new disease-modifying treatments. Genetic association studies have linked epithelial-restricted genes to IPF risk(*4–8*), supporting a conceptual model of IPF pathogenesis wherein the disease develops as a result of repetitive epithelial injury with a dysregulated repair process(*9*, *10*). Single cell transcriptomic studies have revealed extensive changes in cellular phenotypes of, and molecular programs in, epithelial cells in IPF. Also identified was the emergence and persistence of multiple intermediate/transitional states within the distal airway/alveolar niche(*11*, *12*) including KRT5^−^/KRT17^+^/P63^+^ “aberrant basaloid” cells(*13–15*) as prominent disease features. These findings highlight a critical need to better understand the molecular mechanisms which underlie dysregulated epithelial homeostasis and repair in IPF.

The fundamental role of the lung epithelium is to interact with the external environment and maintain the blood-gas barrier. In the alveolar niche, alveolar type 2 (AT2) cells are known to serve as a facultative stem cell, self-renewing and giving rise to new alveolar type 1 (AT1) cells during lung regeneration(*16*). Multiple groups have now identified both homeostatic and disease-emergent cell types/states with distal-airway markers (*SCGB3A2*, *SFTPB*) that may contribute to “attempted” alveolar repair in IPF(*13*) and other chronic lung diseases(*11*, *17*, *18*); these cells share some features with injury-emergent intermediate/transitional states observed in mouse models(*12*, *19–21*). Some of these cells can give rise to mature AT2 cells *in vitro* and likely represent an additional stem-cell reservoir being tapped to regenerate the alveolar niche(*11*, *17*, *22*). The relative contributions of resident AT2 cells and distal airway derived stem cells (i.e. Respiratory Airway Secretory Cells (RASCs)(*17*)/TASCs(*18*)/TR-BSC(*11*)) in repopulating injured alveoli in IPF remains uncertain, yet factors governing the response of each population to repetitive injury are likely key determinants of adaptive versus maladaptive repair.

In the lung epithelium, the Hypoxia-inducible factor (HIF) family of transcription factors has previously been implicated as a determinant of airway epithelial repair following influenza infection(*23*), polarization toward a mucus-secretory cell phenotype(*24*, *25*), and lung endothelial pathology from pulmonary hypertension(*26*, *27*); however a role in lung fibrosis has not previously been established(*28*). HIF signaling can be activated via two distinct but not exclusive mechanisms: 1) impaired degradation via inhibition of prolyl-hydroxylases and ubiquitination when oxygen tension is low, and/or 2) through changes in cell oxidative state (which can be influenced by inflammatory signaling, glycolytic tone and/or other oxidative metabolism/enzymatic activity(*29*)). The tonic degradation of HIFα (both HIF1α and HIF2α [*EPAS1*]) is accomplished through proline hydroxylation and subsequent poly-ubiquitination by Von-Hippel-Lindau (VHL complex). Under activating conditions, HIF1 (or HIF2)α is not degraded, binds HIF1β (ARNT), and is transported to the nucleus where transcription of genes containing a HRE (Hypoxia Response Element) are enhanced. Despite recognizing identical consensus sequences, HIF1α and HIF2α exhibit unique, non-competitive transcriptional regulatory programs(*30*), and also display different activation kinetics(*31*, *32*) with HIF2α demonstrating more prolonged activity with chronic activation.

Using epithelial-targeted Hif deficient mice paired with scRNA-seq, we observed that both AT2 and airway secretory cell contribute to alveolar repair following repetitive injury, and deletion of Hif1/2 in the lung epithelium attenuated fibrosis and promoted cellular programs associated with alveolar repair and maturation. Targeted pharmacologic HIF2 inhibition with PT-2385 (a selective HIF2α: HIF1β dimerization inhibitor) enhanced alveolar repair and differentiation of airway-derived cell populations *in-vivo*. In addition, *ex-vivo* mouse studies showed HIF2-inhibition driven improvement in distal airway to alveolar differentiation. Human distal lung organoid culture revealed enhanced proliferation and importantly suppressed aberrant transitional cell states closely resembling IPF-emergent epithelial cell states. Together, these results indicate HIF2 is a key regulator of distal lung epithelial repair and regeneration, and establish HIF2 inhibition as a novel therapeutic strategy for enhancing lung regeneration.

## Results

To begin investigating the cellular mechanisms that underlie epithelial dysregulation in IPF, we first interrogated single-cell RNA-sequencing (scRNA-seq) data from 67 pulmonary fibrosis (PF) and 49 control (declined donor) lungs (reported in (*33*)) (**Figure 1A-B**). Focusing initially on the IPF-enriched KRT5^−^/KRT17^+^ “aberrant-basaloid” cells, we performed a pathway and transcription factor analysis of genes with Log_2_FC >1. In addition to several pathways that have been previously described in these cells (Epithelial-mesenchymal transition, p53, tumor-necrosis alpha signaling)(**Figure 1C**), transcription factor coexpression analysis revealed significant enrichment of both *HIF1A* (p=4.7×10^−49^) and *EPAS1* (which encodes for HIF2α) (p=2.93×10^−8^) regulated genes, and a “hypoxia module” (comprised of identified HIF regulated genes) was significantly higher in IPF-derived cells compared to controls (**Figure 1D**) (p<2×10^−16^). This suggested that HIF signaling may be a key driver of dysregulated epithelial cell identity and function in PF.

**Figure 1.**
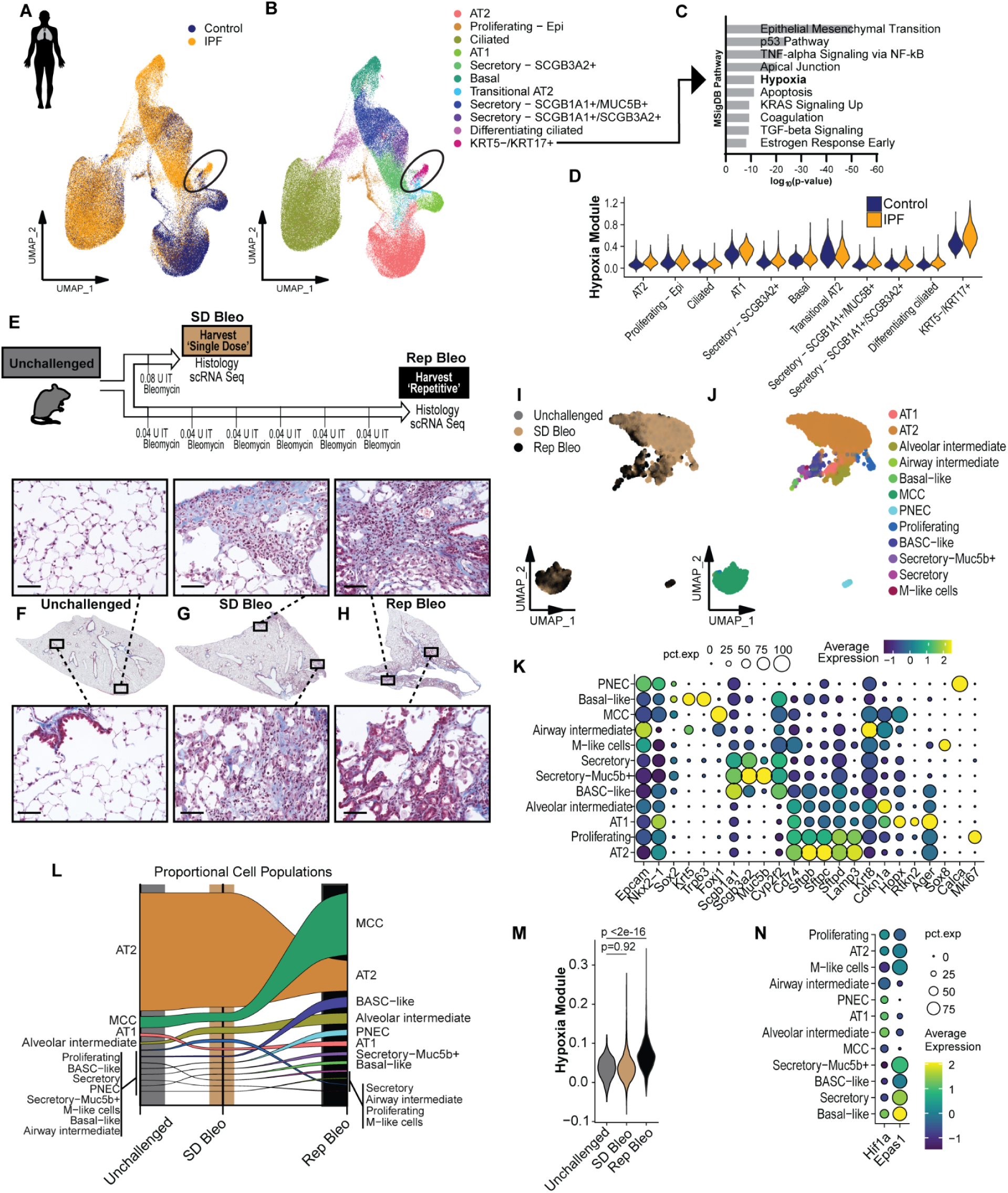
Recurrent injury leads to persistent activation of HIF-regulated programs in the lung epithelium. A-B) UMAP embedding of scRNA-seq of 194,132 epithelial cells from 67 PF and 49 control lungs from reference (*33*) (GSE227136). Circle highlights disease-emergent KRT5^−^/KRT17^+^ cells. C) Pathway analysis of differentially expressed genes with Log_2_-fold change >1 for KRT5^−^/KRT17^+^ cells. D) Hypoxia gene module score (calculated from genes extracted from ARCHs4 transcription factor analysis of HIF1A and EPAS1 of genes in C) with statistically significant increase (p<2×10^16^ pairwise Wilcoxon analysis adjusted for multiple comparisons). E) Schematic of single-dose and repetitive intratracheal (IT) bleomycin mouse models. F-H) Hematoxylin & Eosin stains of representative lung sections from mice following single-dose or repetitive IT bleomycin. Top images demonstrate overall architectural changes. Bottom panels at higher magnification show representative epithelial changes with hyperplasia most prominent in Rep Bleo. Scale bar = 1mm for (a) and 100 µm for (b-c). I-J) UMAP embedding of 6583 cells from unchallenged, single-dose IT bleomycin and repetitive IT bleomycin obtained following FACS-based Cd326 enrichment and 10X 5’ scRNA-seq. These data were jointly analyzed/embedded/annotated with other murine scRNA-seq (**Supplemental Figure 1**) for consistency across figures. K) Dotplot of marker genes driving celltype annotation. L) Alluvial plot showing proportional cell population changes between unchallenged, single dose, and repetitive bleomycin treatment. M) Violin plot of Mouse MSigDB hypoxia module in mice (207 genes) across all epithelial cells from unchallenged, SD Bleo and Rep bleomycin mice. N) Dot-plot of *Hif1* and Hif2(*Epas1*) expression across different cell clusters.

To investigate the mechanisms that underlie resolving vs. nonresolving lung fibrosis in mice, we challenged C57B6 mice with a single-dose of intratracheal (IT) bleomycin (SD Bleo, 0.08 IU) or repetitive biweekly IT bleomycin (Rep Bleo, 0.04 IU) and sacrificed mice 14 days after the final bleomycin dose (**Figure 1E**). Lung histology compared to unchallenged mice (**Figure 1F(a-c)**) showed inflammatory infiltrates (**Figure 1G(a-b)**) and patchy fibrosis (**Figure 1G(c)**) in mice treated with SD Bleo, while mice treated with Rep Bleo exhibited discrete areas of epithelial hyperplasia in the parenchyma (resembling microscopic-honeycombing) (**Figure 1H(a-c)**) along with interalveolar septal thickening and fibrosis. To define the cellular and molecular programs following acute vs. recurrent injury, we then performed scRNA-seq of Cd326+ (EpCAM) sorted epithelial cells (n=3 mice per group). In addition to canonical lineages (AT1, AT2, multiciliated, PNEC[Pulmonary neuroendocrine cells]), we observed a number of injury-emergent cell types/states, including Krt5^+^/Trp63^+^ basal-like cells, Muc5b^+^ secretory cells, airway (Sox2^+^) and alveolar (Krt8^+^/Cdkn1a^+^) intermediate cell populations, bronchiolo-alveolar stem-cell (BASC)-like cells (Scgb1a1^+^/Sftpc^+^)(*34–36*), and Sox8+ microfold-like cells (*37*) (**Figures 1I-K**). Compared to the SD Bleo group, lungs from Rep Bleo mice had expansion of injury-related populations including alveolar intermediate (Krt8^+^/Cdkn1a^+^), BASC-like cells, and Secretory-Muc5b+ cells, as well as significantly reduced AT2 cells (**Figure 1L**). Globally, epithelial cells from mice treated with Rep Bleo showed higher expression of Hif/hypoxia regulated genes compared to unchallenged mice and SD Bleo-treated mice (Mouse Hallmark MsigDB hypoxia gene set, p<2E-16) substantiating hypoxia-driven expression changes unique to the Rep Bleo model (**Figure 1M**). *Epas1* was more highly expressed in key AT2, secretory, and BASC-like populations(*38*)(**Figure 1N**).

Given the potential for at least partial functional redundancy between Hif forms (*32*, *39*), we sought to investigate the role of Hif signaling in chronic epithelial injury by deleting both Hif1 and Hif2(*Epas1*). Lung epithelial-specific Hif1/2 deficient mice (referred to as Hif1/2^Δepi^) (*28*) and cre-negative littermate controls were challenged with Rep Bleo(*28*, *40*) and harvested 2 weeks after the final bleomycin dose (**Figure 2A**). Compared to littermate controls, lungs from Rep Bleo-injured Hif1/2^Δepi^ mice had less architectural remodeling, reduced fibrotic area, smaller lung area occupied by hyperplastic epithelial cells, and lower collagen content (**Figure 2B-E**).

**Figure 2.**
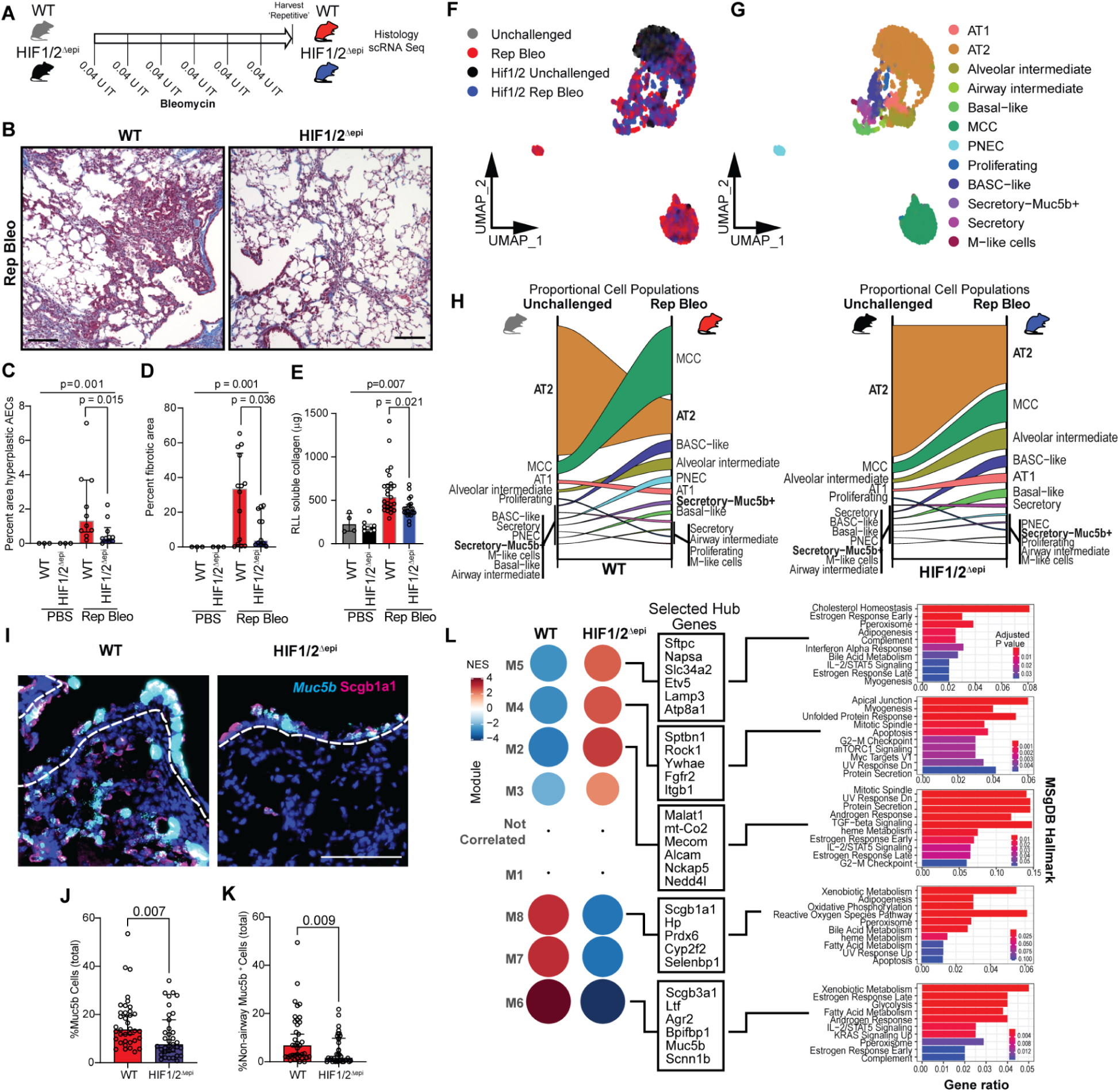
Epithelial deletion of Hif1/2 modulates dynamics from recurrent injury and attenuates experimental fibrosis. A) Dosing and mouse genotype schematic of WT and Hif1/2^Δepi^ mice. B) H&E images of the differential fibrotic response modulated by epithelial Hif1/2 deletion. Scale bar = 1 mm. C) Cell count per 20x field of hyperplastic epithelial cells. Plotted as median ±95% CI. D) Histologic quantification of fibrotic area. Plotted as median ±95% CI. E) Collagen content (Sircol soluble assay) analysis of RLL. Data plotted as median ±95% CI. F) UMAP embedding of 5105 sorted epithelial cells from WT and Hif1/2^Δepi^ mice showing genotype and treatment distribution. Data are pooled from N=3 mice per genotype/treatment condition. G) UMAP of cell type annotation. Cells from Hif1/2^Δepi^ were embedded and jointly annotated (see Supplemental Figure 1). Unchallenged and Rep Bleo WT mice are re-presented for clarity across figures. H) Alluvial plots of cell proportion comparing genotype and response to Rep Bleo. I) RNA-ISH for *Muc5b* and IF of Scgb1a1 from repetitive bleomycin treated Hif1/2^Δepi^ and control mice. J-K) Quantitation of *Muc5b^+^* cells from 10 independent 20x fields across n=4 mice from each genotype. Plotted as median ±95% CI. L) Gene co-expression module enrichment analysis of the airway compartment (BASC, Secretory, Secretory-Muc5b+, Basal), colored by Net enrichment (NES) and plotted by Z-score. Selected genes are presented from hubs within the modules and MSigDB pathway analysis of all module related genes. Additional analysis of module enrichment per cell type and specific Protein-Protein interaction analysis appears in **Supplemental Table 1** and **Supplemental Figure 2–3**).

Next, to investigate the mechanism through which Hif deletion protects against experimental lung fibrosis, we performed scRNA-seq of Cd326+ sorted epithelial cells from unchallenged and Rep Bleo treated Hif1/2^Δepi^ and wild type(WT) mice (**Figure 2F-H**). While there were no compositional differences between unchallenged Hif1/2^Δepi^ mice and WT controls, after Reo Bleo, Hif1/2^Δepi^ had a larger proportion of AT2 cells (38.6% vs 22.3%) and alveolar intermediate cells (13.9 vs 7.9%), and a smaller proportion Secretory-Muc5b^+^ cells (1.0% vs 2.5%) compared to repetitive bleomycin-treated control mice (**Figure 2H**). The increased frequency of distal lung cell types (AT1/AT2/Alveolar intermediate/BASC-like), suppression of PNECs, and rebalancing of the airway populations indicated broad Hif-regulated effects on epithelial population dynamics during repair after recurrent injury. RNA-in situ hybridization/Immunofluorescence for *Muc5b* and Scbg1a1 confirmed attenuation of *Muc5b* expression within(**Figure 2I-J**), and outside of conducting airways (**Figure 2K**), consistent with a direct role of Hif regulation of mucous-secretory cell polarization(*25*).

Focusing on airway-type cells, we then examined gene-coexpression modules to understand differential transcriptional program changes related to Hif-driven signaling. Using CEMiTool(*41*) we identified eight modules, seven of which showed significant genotype-specific enrichment (**Figure 2L**, **Supplementary Data, Supplemental Figure 2**). Modules 2, 4, and 5 (M2/4/5) were most strongly enriched in Hif1/2^Δepi^ mice after Rep Bleo treatment and demonstrate gene co-expression hubs suggesting trans-differentiation of secretory cells toward alveolar fate (*Sftpc*, *Etv5*, *Lamp3*). Notably, M4 shows key hubs associated with 14-3-3 signaling (Ywhae-14-3-3 epsilon) (potentially modulating YAP/TAZ mediated activity(*42*, *43*) and Fgfr2. Module 2 (M2) contains multiple notable hubs associated with airway homeostasis and disease modification (e.g. MECOM in COPD(*44*), Alcam in IPF(*45*)). Further, protein-protein interaction analysis of M2 (enriched with Hif-deletion) identified Wingless (Wnt)-signaling hubs (Glycogen-synthase kinase 3β, Gsk3β) and the transcriptional repressor with the notch pathway Atxn1 (*46*), further supporting the role of Hif-signaling and its interaction with the Wnt/Notch axis with distal lung repair during injury (*23*). This finding specifically positions Hif-activation as an upstream inhibitor of airway to alveolar transition. This is further supported by the enhancement of the AT2-like program within M5 containing protein-protein hubs Cepba and Lrrk2 (**Supplemental Figure 3**). Module 6 alternatively was highly enriched in wild-type (WT) cells and was consistent with airway cell phenotypes, including *Scgb3a1*, *Muc5b*, the transcription factor Ltf and known Hif-responsive *Scnn1b*. This finding is consistent with the other hypoxic-signaling changes noted recently regarding specific secretory populations(*24*, *25*). Pathway analysis of the Hif1/2^Δepi^-enriched modules found associations with proliferation, secretory function, and also biosynthetic machinery associated with cholesterol and lipid synthesis. These data cumulatively suggest that epithelial Hif deletion enables improved adaptive repair through modulation of underlying genomic programs by preventing maladaptive transdifferentiation of mucous secretory cells and facilitation of BASC-like and secretory proliferation programs.

To establish the origin of Hif-activated epithelial cells that emerge following recurrent injury, we performing lineage-tracing studies using *Sftpc-Cre-ER^T2^* (AT2-specific) and *Scgb1a1-Cre-ER^T2^* (secretory cell-specific) mice (**Figure 3A**). After administration of tamoxifen followed by >3 week washout, mice were treated with 4 cycles of IT bleomycin, and CD326^+^/dTomato^+^ cells were sorted by FACS for scRNA-seq (**Figure 3B-C**). Compositional analyses revealed that the Scgb1a1-derived population closely resembles the spectrum of epithelial cell phenotypes recovered following a pan-epithelial sort from mice following chronic injury (MCC, AT2, Alveolar intermediate, and BASC-like cells), suggesting these Scgb1a1-lineage labeled cells were primary contributors to repair of recurrent epithelial injury (**Figure 3D**). Using pimonidazole (Pim) to detect intracellular hypoxia-related reductive signaling (*47*), we observed that in unchallenged mice, there were negligible Pim^+^ cells, whereas abundant Pim^+^ Scgb1a1-traced cells were observed in both the airway and in the repairing parenchyma in Rep Bleo mice, including patches of Spc^+^/Scgb1a1^+^/Pim^+^ cells within areas of parenchymal remodeling, supporting the concept of Hif-driven modulation of repair (**Figure 3E**). Consistent with our initial transcriptomic studies (**Figure 1L**), we identified Hif2α protein stabilization and numerous dTom^+^/Scgb1a1^+^/Hif2α^+^ cells in the lungs of mice following Rep Bleo, while Hif1α^+^ cells were limited to the proximal airways (**Figure 3F**). Together, these data indicate that secretory cells act as prime facultative progenitors in repair of chronic injury, and support the concept that Hif2α activation drives maladaptive repair.

**Figure 3.**
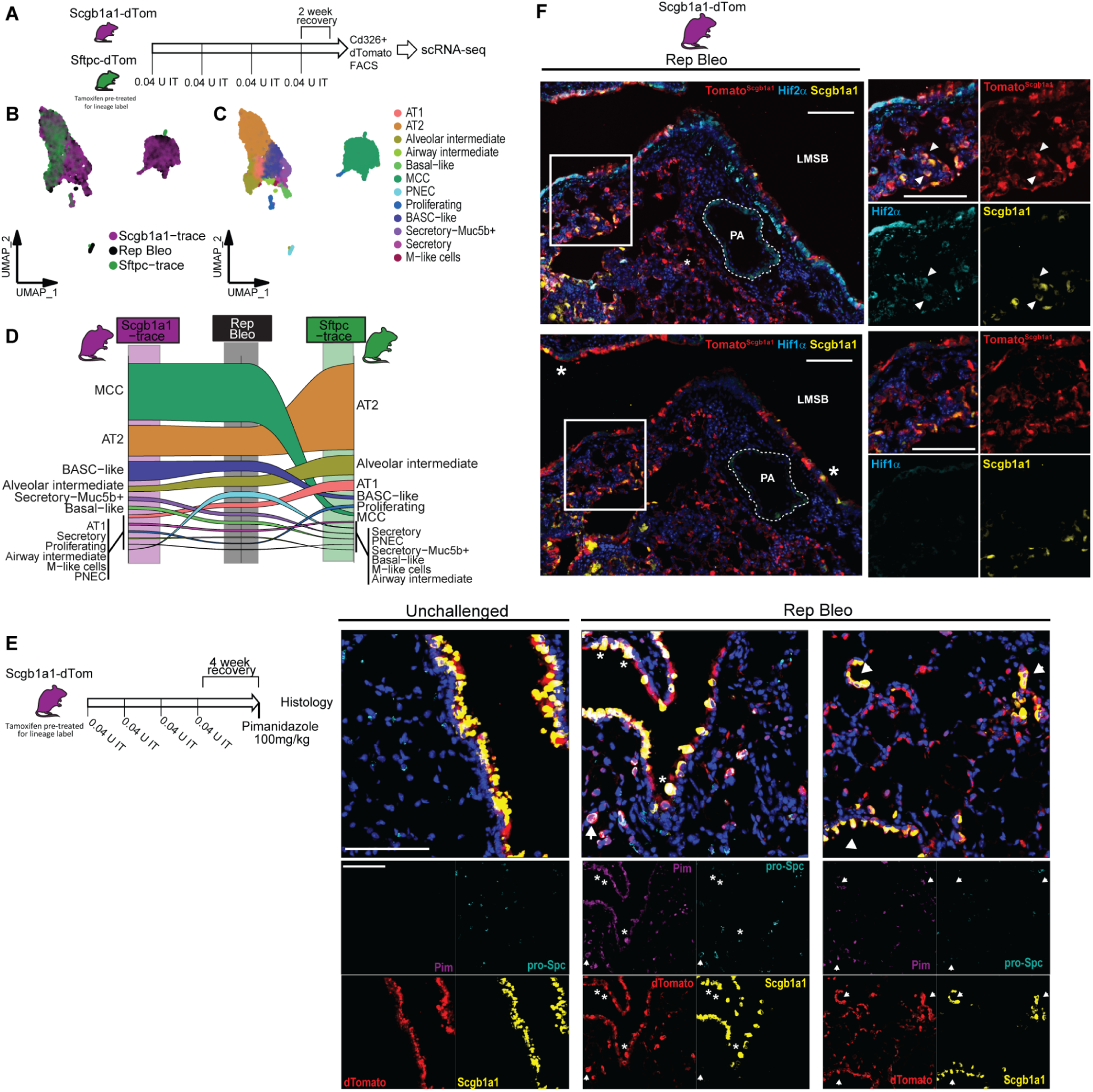
Airway-derived progenitors show Hif2-predominant activation in chronic injury. A) Schematic for lineage-labeled *Sftpc*- or *Scgb1a1*-dTom mice with repetitive bleomycin and subsequent flow-cytometry based dTom+ sorting strategy for scRNA-seq. B-C) UMAP embeddings of 7629 cells by lineage line and by cell type. D) Alluvial plot comparing cell proportion to observed Rep Bleo total unlabeled epithelium compared to dTom^+^ cells from either *Scgb1a1*-traced (left) or *Sftpc*-traced (right). E) Representative Immunofluorescence images of Scgb1a1-traced Rep bleo injured mice treated with Pimonidazole (100 mg/kg) 2 hours prior harvest (detecting intracellular hypoxia or changed redox state) demonstrating extra-airway Pim^+^/dTom^+^ cells both within the airway (stars) and in the parenchyma (arrows) with variable Scgb1a1-protein staining. Of note, there are many instances of Scgb1a1^+^/pro-Spc^+^/dTom^+^/Pim^+^ cells consistent with a cellular state capable of stabilizing Hif-driven signaling. F) Costaining of Scgb1a1-lineage label Scgb1a1 and Hif1 or Hif2 IF using serial sections from Rep Bleo injured mic. White arrows indicated cells with nuclear (active) Hif1 or Hif2. Rare Hif1^+^ airways cells within LMSB. Quantitation of nuclear (active) Hif1 and Hif2 is in Supplemental Figure 4. LMSB = Left mainstem Bronchus, PA = Pulmonary artery. Scale = 100 µm.

Next, we employed a selective and specific small molecule inhibitor (PT-2385) (*48*) which prevents HIF2α:HIF1β dimerization to selectively test the role of Hif2 in regulating alveolar repair during repetitive injury. In these studies, Scgb1a1-lineage labeled mice were challenged with biweekly repetitive IT bleomycin, and PT-2385 (4 mg/kg/day by osmotic pump)(*48*, *49*) was initiated 7 days after the initial dose of bleomycin and continued until sacrifice after 6 cycles of Rep Bleo (**Figure 4A**). Analysis of Hif1 and Hif2 nuclear localization demonstrated PT-2385 appropriately suppressed nuclear localization of Hif2 (**Supplemental Figure 4**). There was a striking increase in dTomato^+^ parenchymal area in PT-2385-treated mice compared to vehicle (DMSO:proplyene glycol)-treated controls (**Figure 4B-C**), with commensurate increased numbers and proportions of parenchymal dTomato^+^ cells expressing the mature alveolar markers pro-Spc (AT2) and/or Ager (AT1) (**Figure 4D-F**). In addition, while *Muc5b* expression remained unchanged in the airways, there were significantly fewer *Muc5b*^+^ cells outside of conducting airways (**Figure 4G-I**), similar to our observations in mice with combined Hif1/2 genetic deletion (**Figure 2J-K**). These data collectively support the mechanistic concept that interrupting persistent Hif2 activation in the regenerating lung epithelium can enhance distal lung epithelial and alveolar repair.

**Figure 4.**
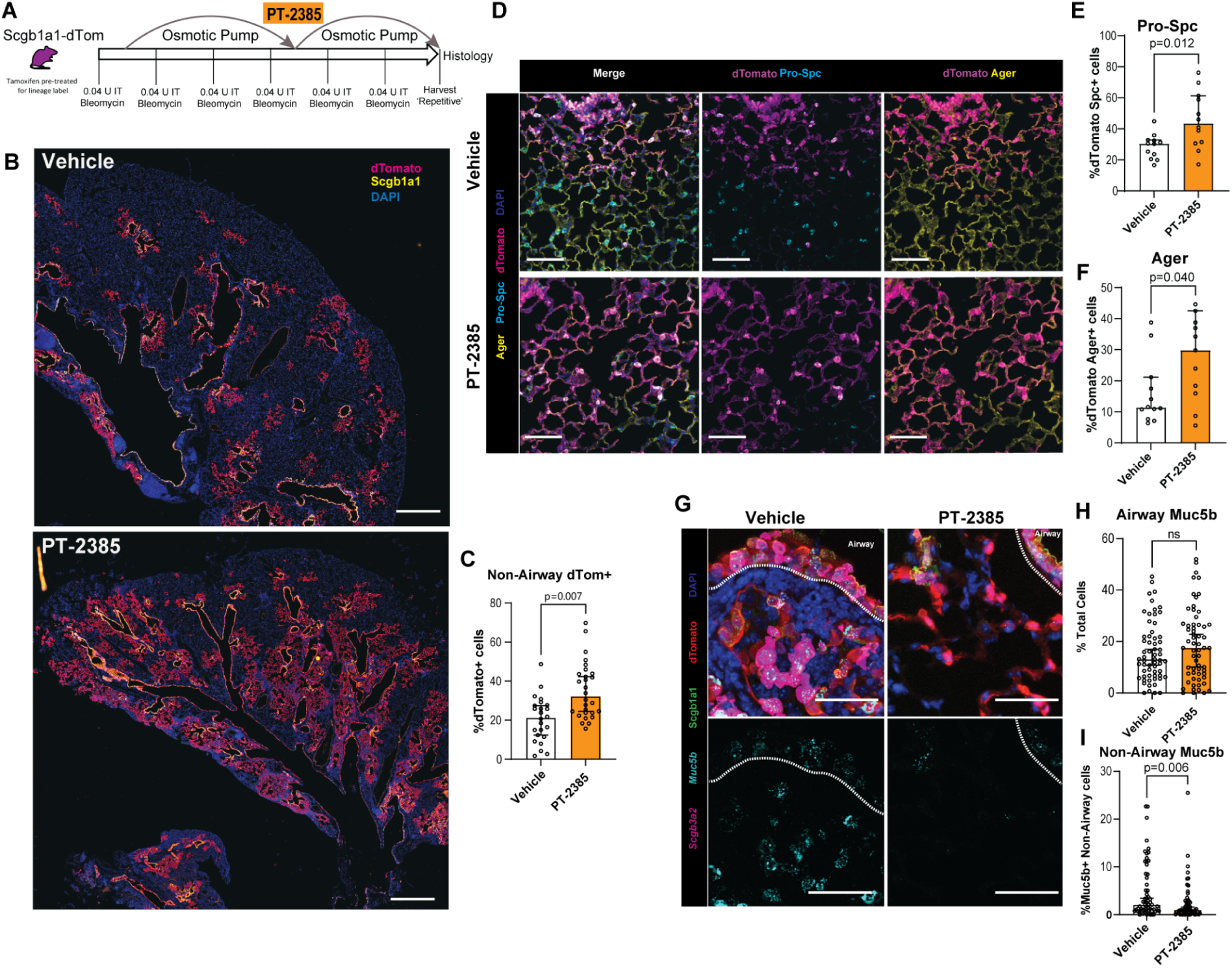
Pharmacologic Hif2-inhibition enhances airway-derived adaptive epithelial repair following recurrent injury. A) Schematic of Rep bleo in Scgb1a1 lineage labeled mice with bi-weekely IT bleomycin. The Osmotic pumps containing PT-2385 (propylene glycol/DMSO vehicle) were implanted subcutaneously 1 week following the initial dose of bleomycin and were then replaced with fresh pumps following the forth bleomycin administration. Mice were harvested for analysis two weeks following the sixth IT bleomycin dose. B) Stitched images of Scgb1a1-lineage labeled repair in Vehicle vs PT-2385 in repetitive bleomycin. Scale bar = 1mm. C) Quantitation of dTomato^+^ Scgb1a1-lineage derived cells outside airways. N=6 mice per group with at least 4 sections per mouse quantified. Plotted as median ±95% CI. D) IF of pro-Spc, Ager, and Scgb1a1-lineage labeled dTomato. Two 5×5 stitched images were quantified per mouse with n=6. Plotted as median ±95% CI. Scale bar = 50 µm. E-F) Quantitation of pro-Spc or Ager with dTomato dual positivity. Two 5×5 stitched images were quantified per mouse comprising between 7900 to 15000 cells were quantified. n=6. Plotted as median ±95% CI. For F, Welch’s test was used in accordance with a non-significant variance difference. G) Combined RNA-ISH (*Muc5b* and *Scgb3a2*) and IF (Scgb1a1, dTomato). Dotted line delineates the airway from adjacent parenchymal structures. Scale bar = 50 µm. H-I) 10 separate images from N=6 mice were masked to delineate airway (H) from parenchyma (I) and analyzed.

To determine whether the effects of Hif2 activation were due to autonomous effects on regenerating epithelial cells, we sorted Cd326^+^/dTomato^+^ cells from naive Scgb1a1^dTom^ mice and established feeder-free alveolar organoids using “serum-free, feeder-free media” (SFFFM)(*50*) in matrigel droplets (**Figure 5A-B**). Consistent with our *in-vivo* findings, PT-2385-treated Scgb1a1^dTom^ derived organoids had more robust AT2-marker acquisition (proSPC^+^) compared to vehicle-treated controls (**Figure 5C-D**, while maintaining AT1 differentiation potential (**Fig. 5E**, **Supplemental Figure 5**). These indicate that Hif2-activation in airway-based repairing epithelium restrains alveolar differentiation potential, which can be rescued through Hif2-specific inhibition.

**Figure 5.**
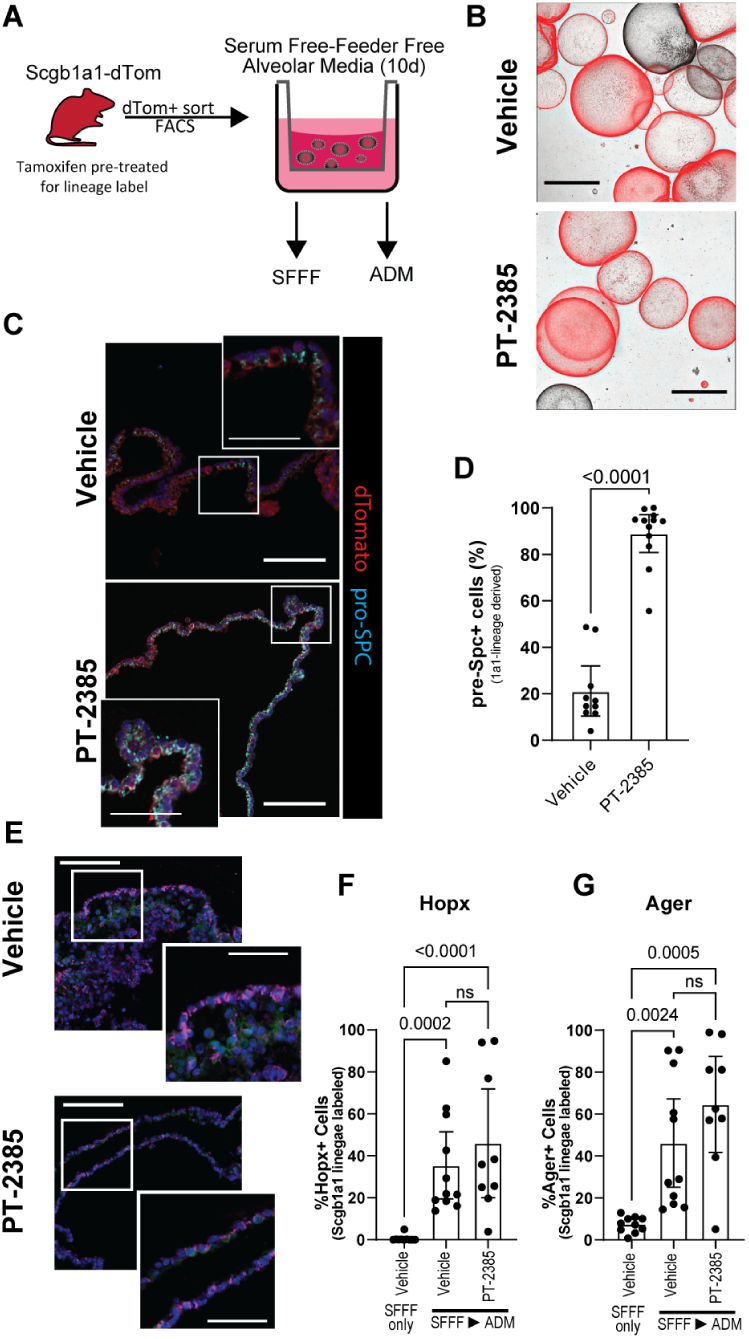
Inhibiting Hif2 promotes alveolar fate of airway-derived progenitors. A) Schematic of FACS sorting of *Scgb1a1*-lineage labeled cells for subsequent organoid culture derived from pooled cells from three mice. B) Stitched overlay projection of brightfield and dTomato fluorescence of SFFF organoid outgrowth after 10 days. Scale bar = 1mm. C) IF of organoids for pro-Spc and dTom^+^ cells. Scale bar = 100 µm, inset scale bar 50 µm. D) Quantitation of pro-Spc^+^ Scgb1a1-lineage labeled cells per field of organoid, n=10 organoids per group across three technical replicate wells. E) AT1-marker staining of Scgb1a1-derived organoids after 7 days of SFFFM expansion followed by transition to ADM for an additional 6 days. Scale bar = 100 µm, inset scale 50 µm. F-G) Quantitation of Hopx and Ager IF. Unique fields (comprising different organoids) spanning 2 technical replicates are displayed.

We then turned to a human alveolar organoid model to establish whether HIF2 also regulates alveolar progenitor potential in the human lung (**Figure 6A**, **Supplemental Figure 6**). Following initial expansion after isolation of CD326+ epithelial cells from declined donors under serum-free feeder-free (SFFF) conditions (*11*, *50*), we disaggregated and re-established feeder-free organoids that were immediately treated with 20µM PT-2385 or vehicle (DMSO). By day 14, PT-2385 treated organoids were more abundant (**Figure 6B-C**) and demonstrated robust lysotracker staining (indicating lysosomal related organelles associated with surfactant production)(*51*), results we confirmed using primary AECs from IPF lungs (**Supplemental Figure 7**). Intriguingly, pro-SPC, SCGB3A2 and KRT8^+^ cells could be found within the same organoid, suggesting some plasticity among these cell types under these conditions (**Supplemental Figure 6C**).

**Figure 6.**
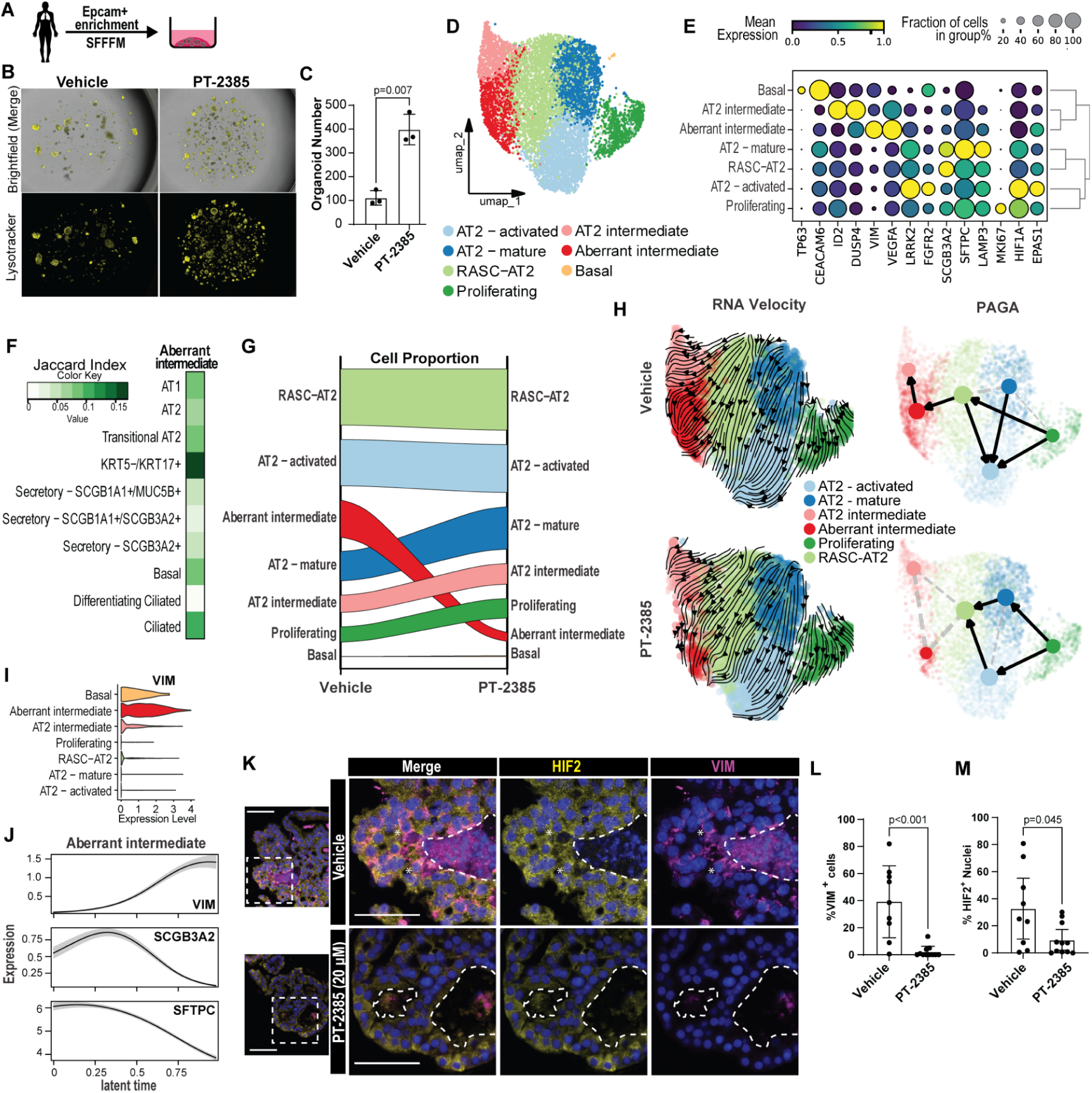
Inhibiting HIF2 enhances human alveolar organoid growth and prevents the emergence of aberrant intermediate cells. A) Human Organoid generation using declined donor tissue or ILD explants via CD326 enrichment and culture in SFFFM. B) Brightfield and Lysotracker images of Vehicle and PT-2385-treated organoids (passage 1). Scale Bar = 1mm. C) Quantitation of outgrowth from vehicle and PT-2385-treated organoids at 14d of passage 1. Analyzed with Welch’s t-test after ensuring variance did not significantly differ. D) UMAP embedding of cell cluster annotation for 8694 cells across Vehicle (DMSO) and PT-2385 treatments. Treatment group embeddings appear in **Supplemental Figure 6**. E) Dotplot for cluster annotation and marker gene expression. F) Jaccard analysis of cluster-specific DE-genes (Log2FC>0.5 and p<0.001) between the Aberrant intermediate cluster and full lung epithelial dataset appearing in Figure 1A. G) Alluvial plot comparing cell proportions between Vehicle or PT-2385 (20 µM) - treated organoids. H) RNA velocity stream embeddings and PAGA representations for trajectory analysis of separately analyzed Vehicle and PT-2385 organoids demonstrating different dynamics between clusters, particularly within the RASC-AT2 cluster based on HIF2 inhibition. Further latent time embedding, differentially predicted terminal states and absorption probabilities appear in Supplemental Figure 8. I) Violin plot of *Vimentin* (*VIM*) expression across cell-types. J) Modeled marker gene expression along Cellrank-estimated latent time trajectory for Aberrant intermediate cells demonstrating predicted progression through RASC-AT2 state. K) Representative IF of isolated organoids stained for Vimentin, HIF2α and their colocalization. Dotted lines delineate internal cavities of organoids for context. Stars indicate cells with both HIF2a nuclear localization and cytoplasmic Vimentin. Scale bar = 50 µm. L-M) Quantification of Nuclear HIF2α or Vimentin. 5 different organoids quantified across two technical replicates. Plotted as median ±95% CI.

We performed scRNA-seq of Vehicle and PT-2385 organoids which demonstrated seven cell-types/states, including AT2, RASC-AT2 (*SCGB3A2*^+^/*SFTPC*^+^), along with other activated/intermediate states (**Supplemental Figure 6**). A unique population of “Aberrant intermediate” cells was much less frequently observed in PT-2385 treated organoids (18.4% vs 4.4%) (**Figure 6D-G**, **Supplemental Figure 6**). Jaccard analysis comparing the Aberrant Intermediate organoid population and cell-populations observed in the IPF lung epithelium (from **Figure 1A-B**) demonstrated the greatest degree of similarity (p = 2×10^−174^) was with the highly disease-enriched (and hypoxic-module expressing) KRT5^−^/KRT17^+^ cells (**Figure 6F**). Pathway analysis of these aberrant intermediate cells suggested both NF-kB activation (associated with epithelial-mesenchymal-transition (EMT) and maladaptive repair(*52*)) as well as hypoxic signatures, with upregulation of key HIF-responsive genes including *VEGFA*, *ATF3*, *TXNIP* (**Supplemental Figure 6**).

Several recent reports have implicated IPF related basal cell populations with aberrant phenotypes and fibrotic responses (*53*, *54*), however the transcriptional similarity of an analogous disease emergent populations from distal lung organoid culture would support the possibility that facultative stem cell populations (RASC-AT2 or AT2) as possible sources for these cell types *in-vivo*. mRNA-splicing-based trajectory analysis demonstrated that HIF2 inhibition led to differential dynamics, particularly between the RASC-AT2 and Aberrant intermediate clusters. Partition-based graph abstraction (PAGA)-based fate prediction suggested the RASC-AT2 to Aberrant intermediate transition was diminished by HIF2 inhibition, while the proliferating cell to AT2 mature and then RASC-AT2 was enhanced (**Figure 6H**). Further analysis using velocity kernel based estimators predicted different terminal states and thus different quantitative dynamics for the RASC-AT2 population (**Supplemental Figure 8**). We observed that aberrant intermediate cells expressed high levels of Vimentin (*VIM*) (a marker of EMT (*55*) and a known HIF-responsive gene (*56*)) both in terms of average expression and when modeled over the calculated Aberrant intermediate trajectory (**Figure 6I-J**), similar to that observed in KRT5^−^/KRT17^+^ in IPF(*13*, *14*). Immunofluorescence staining revealed frequent VIM^+^ cells with nuclear (i.e. active) HIF2 in vehicle-treated organoids, which was highly suppressed by PT-2385 (**Figure 6K-M**). AT1 marker acquisition following addition of differentiation media was unchanged (**Supplemental Figure 9**). These data collectively demonstrate that in a human distal lung organoid model, HIF2 activity regulates the phenotypic plasticity of distal lung epithelial cells, and while HIF2 activation promotes adoption of an aberrant intermediate cell state, this can be effectively modulated via small-molecule-based inhibition to promote functional repair and maturation.

## Discussion

While the lung epithelium exhibits remarkable ability to repair and regenerate after acute injury, recurrent and/or chronic insults lead to enduring changes in the composition and molecular programs active in the lung epithelium, resulting in pathologic structural remodeling and progressive decline in lung function. In this study, we found that persistent activation of HIF-signaling in repair-associated cell types/states is a hallmark of dysfunctional epithelial repair in the IPF lung epithelium and experimental models of recurrent lung epithelial injury. Further, these dysfunctional repair-associated cells arise primarily from airway epithelial progenitors and exhibit specific activation of HIF2. Targeted inhibition of HIF2 promoted adaptive alveolar epithelial repair *in vivo* and prevented emergence of ectopic mucous secretory and aberrant basal-like cells. Together, these findings indicate that HIF2-inhibition is a promising and novel therapeutic strategy to enhance lung repair in IPF and other chronic lung diseases.

Although there has been extensive study of facultative progenitor cell populations that participate in alveolar repair following severe acute lung injury(*16*, *17*, *57–66*), there has been limited investigation of cellular and molecular mechanisms that regulate alveolar responses to recurrent or chronic injury. We found that recurrent epithelial injury results in mobilization of distal-airway derived progenitors, and that distal-airway-derived regeneration-associated intermediate/transitional cells with persistent activation of Hif/hypoxia related signaling failed to mature into functional alveolar epithelial cells. In this setting, we observed that selective inhibition of Hif2 in these regenerating airway-derived epithelial cells was sufficient to redirect their fate away from airway/mucous-secretory-lineages and promote alveolar fate acquisition and maturation to mediate functional lung repair. While there have been multiple characterizations of airway organoids differentiating AT2-like Sftpc-expressing cells (*63*, *64*, *67*, *68*), we observed that Hif2 inhibition dramatically enhanced this cellular transition.

Hif-related signaling, primarily Hif1α, has been studied in the context of acute lung injury. In the acute setting following alveolar injury, several studies suggest Hif1α plays a protective role in AT2 cells(*69–71*). In contrast, deletion of Hif1α in Sox2^+^ airway-derived progenitors following severe influenza injury attenuated lung injury and the development of metaplastic Krt5^+^ “pods” via regulation of the Notch pathway(*23*). One potential unifying explanation for these observations is that Hif1α plays distinct roles in specific cellular contexts, and that differences in Hif1α kinetics influence adaptive vs. pathologic effects. There has not been previous study of Hif2 specifically in these contexts, and while these data indicate that Hif2 activity restrains progenitor function and alveolar fate potential while promoting airway-type metaplasia, it remains possible that Hif1 plays an additional independent or niche regulated role. Further, it is also possible that requirements for facultative stemness of AT2 cells (i.e. biosynthetic burden for lipid synthesis) could also be modulated through metabolic changes driven by HIF2, or that HIF2 influences sensitivity to AT2 differentiation cues via modulation of the WNT/NOTCH balance (*72*).

Hypoxia and HIF-signaling have been previously been linked to mucous secretory cell differentiation (*24*, *73*), and in this study we found that either genetic deletion of Hif1/2 or small-molecule-based inhibition of Hif2 was sufficient to attenuate mucous-secretory cell differentiation. As expansion of mucous secretory cells is a prominent feature of the IPF lung (*13*) and may be driven at least in part by a promoter polymorphism in the gene encoding for MUC5B (*4*, *33*), this work builds upon other recent work which indicates that targeting HIF may be an effective means to ameliorate the untoward effects of mucous hypersecretion in chronic respiratory diseases while providing a tenable therapeutic target as a disease modifying agent(*25*).

In addition to increased mucous secretory cells, one of the most striking findings of recent single-cell transcriptomic studies has been the identification of KRT5^−^/KRT17^+^ “aberrant basaloid” cells in IPF lungs(*13*, *14*). While questions remain as to the origin (*13*, *53*, *74*, *75*) and specific functions of these cells, a role in amplifying fibrotic signaling seems likely. We were surprised to find that “Aberrant intermediate” transcriptionally resembling some features of “aberrant basaloid” cells emerged in feeder-free culture of human AECs (despite TGFβ inhibition), and our trajectory analyses suggested they arise from *SCG3A2*^+^ “RASC-like” cells in this model; further, their emergence could be largely prevented by HIF2-inhibition. These results suggest that HIF2-signaling, independent of effects on the TGFβ pathway, promotes the emergence and persistence of intermediate/transitional cell states and prevents AEC maturation.

While together these data provide substantial evidence that specific activation of HIF2 is a hallmark of chronic lung epithelial injury and can be effectively targeted to promote lung repair, a number of additional questions remain. Despite some evidence of cellular hypoxia in areas of Hif-activation, it is possible that non-hypoxic mechanisms of HIF activation are as or more important regulators of observed HIF activity in these contexts; the distinct contributions of hypoxic vs “pseudo-hypoxic” activation of HIF1 vs. HIF2 in distinct spatial niches remains to be determined. Further, it is also possible that indirect effects of HIF forms on each other via reciprocal co-regulation (*29*, *76*) contribute to the observed effects on epithelial repair. These studies focused specifically on the role of HIFs in the lung epithelium, but available data suggest this family of transcription factors may also play an important role in fibroblasts(*77–79*), macrophages(*80*, *81*) in addition to well established roles in the endothelium. Finally, although we have attempted to maximally explore HIF2 inhibition in the best available preclinical models for IPF, measurement of clinical endpoints and further human studies will be required to understand the full potential benefit from targeting this pathway. Moreover, while use of PT-2385 appears promising, substantial additional work is needed to optimize *in-vivo* systemic or topical dosing strategies.

In summary, recurrent injury to the distal lung/alveolar epithelium leads to persistent activation of HIF-regulated signaling in airway-derived progenitors which promotes dysfunctional epithelial remodeling. Selective inhibition of HIF2 is a novel therapeutic strategy which has the potential to enhance functional alveolar repair and preserve or enhance lung function for patients with IPF and other chronic lung diseases.

## Methods

### Study design

*Research Objectives*: The overarching objective of the study was to interrogate mechanisms that regulate alveolar repair of recurrent injury. *Research units of investigation* included primary cells/samples from pulmonary fibrosis and control lungs, transgenic mice, and primary cells in culture. *Experimental Design*: The experimental design was a controlled laboratory study including genomic data analysis, mouse models, and organoid studies. *In vitro* work was performed in a minimum of three technical replicates per experimental condition and always jointly analyzed where indicated. For image analysis, quantitation was performed using the same cell identification and size dimension, channel positivity thresholds, and anatomical landmarks through the use of automated image analysis software (Halo, Indica Labs) established based on each immunofluorescent panel with appropriate unstained and secondary-only negative controls. Experimental and Control samples were stained from the same stock solutions and acquired under identical settings. Male and female mice were used in all experiments in as close to equivalent proportions as possible. Statistical methods are discussed in detail below; broadly non-parametric analyses were used and adjusted for multiple comparisons where appropriate. No outliers were excluded from analysis. Animals were only excluded if there was pump loss as detailed below, otherwise all were used in analysis. For human primary-cell studies derived from declined donor samples or post-lung transplant ILD explants, cells from both adult male and female donors were used as they became available through the duration of the work. *Blinding*: Wherever possible, data were generated and/or analyzed blinding to treatment group/genotype, use of automated and thresholded image analysis in bulk further supported impartiality.

### Human samples and subjects

Human lung tissue samples used for scRNA-seq and/or organoid cultures were obtained from PF lung removed at the time of explant and declined donor (control) lungs as previously described (Vanderbilt IRB #’s 060165, 192004).

### scRNA-seq reanalysis

Integrated epithelial cell scRNA-seq data from GSE227136(*33*) were reanalyzed to specifically assess the distal airway and alveolar compartment. A subset of the overall object containing data from IPF and control patients was used to perform find differentially expressed genes in disease-emergent KRT5^−^/KRT17^+^ “aberrant basaloid” cells compared to AT1, AT2, Transitional AT2, Secretory – SCGB3A2+ cells using the FindMarkers function in Seurat with a ‘negbinom’ test. All genes with a Log_2_FC>1 were analyzed for pathway enrichment with MSigDB and with ARCHs4 transcription factor coexpression. ARCHS4 analysis found HIF1A and EPAS1 to be significantly enriched, so a hypoxia module was constructed from the 90 genes represented in the HIF1A set. The module was applied to the entire dataset using default AddModuleScore with default parameters within Seurat and plotted by disease status. Statistical comparisons were made using pairwise Wilcoxon Rank-Sum tests.

### Animal studies

Eight to twelve-week old male and female mice (C567Bl6 background) were used for experiments. *S*ftpc*^tm1(cre/ERT2)Blh^* (*Sftpc-CreER*), *Scgb1a1-CreER^tm1(cre/ERT2)Blh^ (Scgb1a1-CreER)*, and *Rosa26R-CAG-lsl-tdTomato* (tom) were purchased from Jackson Laboratories (# 028054, 016225, 007914). Mice were housed in the animal care facility at Vanderbilt University Medical Center and allowed ad-libitum access to food and water. Epithelial Hif1/2 deficient (Hif1/2^Δepi^) were generated as previously described (*28*) by crossing constitutive Sftpc-Cre mice (*84*) with Hif1/Epas1(Hif2) floxed animals. Cre-negative littermate animals were used as controls for all studies. All studies were approved by the Institutional Animal Care and Use Committee (IACUC) at Vanderbilt.

### Bleomycin fibrosis models

Intratracheal bleomycin (obtained from Vanderbilt University Hospital Pharmacy) (0.04 IU per dose/100uL/mouse/instillation, suspended in sterile PBS) was administered by intratracheal instillation in 4-6 biweekly doses as previously described (*28*, *40*) or in a single 0.08U/100uL/mouse/instillation. Sterile saline in equivalent volume was used as control.

### Lineage-tracing

For lineage tracing and cell-sorting experiments, *Sftpc*-*CreER*^tom^ and *Scgb1a1-CreER*^tom^ mice were administered tamoxifen chow (tamoxifen citrate, Envigo TD.130860, 400 mg/kg diet) x 5 days followed by washout of at least 21 days then used in indicated experiments.

### PT-2385 *in vivo* administration

PT-2385 (MedChemExpress; Cat. No.: HY-12867) was solubilized in a 1:1 v/v admixture of Propylene glycol (Sigma Aldrich) (sterile filtered before use) and DMSO (after compound first being dissolved in DMSO component) to form a colorless solution. This solvent mixture was devised due to the need for high compound concentration and pump bladder compatibility. Under sterile conditions, Alzet osmotic pumps (Model 2004) were loaded per manufacturer instructions with PT-2385 or vehicle alone then primed overnight at 37C in PBS per manufacturer instructions before implantation. Under IACUC-approved surgical conditions, Scgb1a1^tom^ mice were anesthetized using isoflurane and after hair removal and topical sterilization a small incision was made dorsally, anterior the scapulae, and using blunt dissection a pocket was created posteriorly. The pump was then inserted into the pocket and the incision was closed using suture. Wounds were checked daily for the first week then every 48 hrs thereafter. Pumps were removed/reimplanted at the indicated time after additional doses using the same procedure. In the event of pump loss, mice were removed from the experiment.

### Generation of single-cell suspensions

*Mouse:* Following cardiac puncture and perfusion with sterile PBS, lung tissue was isolated to the level of the main carina and rinsed 1x in sterile PBS then placed into a digest mixture in C-tubes (Miltenyi): phenol-free DMEM [high glucose + 25mM HEPES] (2.5 ml/lung up to 10 ml total per C-Tube) with 1mg/ml Type 1 Collagenase (Sigma Aldrich; catalog no. SCR103), 1U/ml Dispase (Roche; catalog no. 04942078001), Aprotinin (2µg/ml), Pepstatin (1 µg/ml), Leupeptin (1 µg/ml) (all RPI, catalog numbers A20550, L22035, P30100) and PMSF ([Phenylmethylsulfonyl fluoride] 1mM final, RPI P20270; added immediately prior to adding lung tissues to digest solution). Tissue was then digested and disaggregated using GentleMacs cells dissociator. Following heated run on the GentleMACs, digests were quenched with an equal volume of phenol-free DMEM with 25mM HEPES + 6mM EDTA (to quench cation dependent proteases and assist cell dissociation) and ensuing cell suspensions were sequentially passed through 100 and 40 μm filters to obtain single cell suspensions which were counted then used in downstream processes.

*Human*: lung tissue was obtained after no more than 12hrs storage at 4C and using appropriate BSL-2 precautions, dissected and distal lung obtained and diced into ~5mm cubes then mashed. 4g of tissue/10ml digest solution [phenol-free DMEM [high glucose + 25mM HEPES] (10ml total per C-Tube) with 1mg/ml Type 1 Collagenase (Sigma Aldrich; catalog no. SCR103), 1U/ml Dispase (Roche; catalog no. 04942078001), Aprotinin (2µg/ml), Pepstatin (1 µg/ml), Leupeptin (1 µg/ml) (all RPI, catalog numbers A20550, L22035, P30100) and PMSF ([Phenylmethylsulfonyl fluoride] 1mM final, RPI P20270; added immediately prior to adding lung tissues to digest solution]. Tissue was then digested and disaggregated using GentleMacs cells dissociator. Following heated run on the GentleMACs, digests were quenched with an equal volume of phenol-free DMEM with 25mM HEPES + 6mM EDTA (to quench cation dependent proteases and assist cell dissociation) and ensuing cell suspensions were sequentially passed through sterile woven gauze, 100 and 70 μm filters to obtain single cell suspensions which were counted then used in downstream processes.

### FACS-based experiments

For FACS, the buffer used for antibody binding and washes composed of phenol-free DMEM with 25mM HEPES/3mM EDTA/0.1% BSA which was 0.2 µm vacuum filtered prior to use. Cells were suspended at 3×10^6^ cell/ml and incubated for 30 min at 4C with 1:100 dilutions of the following antibodies: [APC dump channel: Ter-119 (Biolegend cat no. 116212), CD45 (Biolegend cat no. 103112), CD31 (Biolegend cat no. 102409)] and anti-Cd326 PE-Cy7 (Biolegend cat no.118216) along with 1.25µM Calcein Violet AM (Invitrogen catalog no. 65-0853-78). Following incubation, cells were washed with a 10x volume of buffer (i.e. 3ml for 300µl incubation volume), pelleted at 400g for 5 minutes, then resuspended at 1×10^6^ cells/ml for FACS sorting on a BD 4-laser FACSAria III within a Baker Bioprotect IV hood. Samples were gated as CvAM^+^/ Cd45^−^/Cd31^−^/Ter-119^−^/Cd326^+^ using a 70 µm nozzle. A minimum of 30,000 cells were collected and directly taken for scRNA-seq processing.

### Column enrichment

For column-based magnetic bead isolation, buffer used for antibody binding, column washes and elution was composed of phenol-free DMEM with 25mM HEPES/3mM EDTA/0.1% BSA which was 0.2 µ vacuum filtered prior to use. After sieving, cells were counted and resuspended at 1×10^7^ cells/90µl and 20 µl/1×10^7^ cells of freshly vortexed CD326 microbeads (Miltenyi, catalog no. 130-061-101) were added and incubated on ice with agitation every 5 minutes for 15 minutes. Cells were then washed with 5ml FACS buffer, re-pelleted, and resuspended in 500 µl for application to LS magnetic bead column (Miltenyi, catalog no 130-042-401) on a QuadroMACS (Miltenyi) magnetic separator. Once bound, columns were washed with 12 ml of FACS buffer prior elution. Cells were pelleted and resuspended in appropriate buffer/media for downstream applications.

### scRNA-sequencing

Single-cell RNA-sequencing was performed using the 10X Genomics Chromium 5’ assay targeting capture of 10,000 cells per lane, followed by library sequencing on an Illumina Hiseq4000 or Novaseq6000 targeting 50,000 reads/cell. Alignment and demultiplexing was performed using CellRanger v7.0 aligning to the appropriate reference genome (mm10-2020 for mice, hg38-2020a for human). Murine data was post-processed using Cellbender(*85*), and filtered_output counts matrices were used for downstream analysis.

### scRNA-seq analysis

Seurat v4.3.0 was used to perform dimensionality reduction, clustering, and visualization for the scRNA-seq data. Individual sample output files from CellRanger Count were read into Seurat to generate a unique molecular identifier count matrix that was used to create a Seurat object containing a count matrix and analysis. All Murine Seurat objects were combined into a merged dataset (**Supplemental Figure 1**). The same process was followed for human data. Following combined object generation, cells containing between 500-6000 identified genes and <10% mitochondrial reads were retained. SCTransform with default parameters including mitochondrial and ribosomal genes. Harmony(*86*) was used to integrate the murine dataset prior PC analysis, clustering, and UMAP generation. Major celltype clusters were iteratively annotated using major lineage markers, doublets manually removed, and ultimately subclustered to obtain the final cell type annotations.

### Differential Expression Analysis

Differentially expressed genes between cell types were calculated using the negative binomial model as implemented in the Seurat FindMarkers function with logfc threshold of 0.25.

### CemiTool analysis

The R package CemiTool(*41*) was used to perform modular co-expression analysis on the secretory and basal cell clusters. Briefly, objects containing expression counts and cell genotype or cell cluster identities were extracted and run within Cemitool. Threshold value was assigned following assessment of r^2^ scale free topology and mean connectivity outputs. The following input was used to run the analysis: cemitool(counts.ko.sec, sec_annot_for_KO, gmt = reactome_gmt, interactions = interactions, set_beta = 5, apply_vst = T, filter_pval = 0.15, cor_method = ‘spearman’, cor_function = ‘bicor’, gsea_max_size = 1500, plot = T, plot_diagnostics = T, verbose = T). The ‘bicor’ function was used to better handle correlation analysis given the missingness associated with scRNA-seq data. Murine Protein-Protein interactions were obtained from HitPredict v4(*82*).

### Jaccard Analysis

Systematic assessment of gene representation similarity was performed using the GeneOverlap R package (https://github.com/shenlab-sinai/GeneOverlap) on differentially expressed genes from the GSE227136 Human Epithelial object (Figure 1A) and the Aberrant intermediate cluster on all genes with an adjusted p value <0.001 and log_2_FC >0.5. p values for Jaccard significance were calculated using a gene library of 23749 total genes.

### Trajectory Analysis

Trajectory inference was performed using a combination of scVelo and CellRank. Briefly, spliced/unspliced loom files were generated using velocyto(*87*) and merged with objects obtained above following QC and cluster assignment. The merged objects were then processed with scVelo(*88*) with default parameters using a dynamical model including calculation of latent time. PAGA with velocity directed edges were plotted from this data. Next, using CellRank(*89*), a velocity kernel was calculated using monte carlo approximations to obtain the streamed velocity embeddings for confirmation. Terminal states and absorption probabilities were calculated using a Generalized Perron Cluster Cluster Analysis (GPCCA) estimator with 4 fit states based on assessment of a Schur decomposition to ensure appropriate fitting and default determination of macro and terminal states within the estimator function. These data appear in **Supplemental Figure 8**.

### Immunohistochemistry and Immunofluorescence

#### Mouse Tissue

Following cardiac puncture and perfusion with sterile PBS, Lungs were insufflated with neutral buffered formalin (Fischer Scientific) at 25cmH_2_O then fixed at 4C at least overnight. Further dissection was then performed if needed, and lungs were loaded into cassettes and processed and paraffin embedded. 5 μm sections were used for all histological stains.

#### Organoids

At the indicated time points, organoid-containing matrigel droplets were removed from the incubator and washed 1x 5 min with prewarmed sterile PBS followed by fixation with 4% paraformaldehyde overnight at 4C. Fixed droplets were then washed with PBS and stored until pre-processing. In order to maintain organoid integrity, whole droplets were gently removed from underlying culture substrate using a specially created #16 Blade scalpel with a 90 degree bend introduced half way down the blade to allow easy removal of the whole droplet. Intact droplets were then moved to 2 ml Eppendorf tubes (enabling consolidation of replicates if needed) via cut-off 1ml pipette tips precoated with 1% BSA (to minimize sticking) spun at 100g for 2 minutes for gentle pelleting, resulting supernatant removed, and Histogel (Thermo Scientific) (equivalent in volume to original matrigel, i.e 3x 50 μL droplets would have 150 μL Histogel) was added to resuspend whole fixed droplets. The histogel plugs were then placed on ice to harden then removed using a small blunt balance scoop and placed into a cassette for processing.

#### General Protocol

Slides were heated (30 minutes at 60°C) then washed twice in xylene for 5 minutes, and sequentially rehydrated in ethanol (5min in 100%, 95%, 70%, 50%). Paraffin sections were then antigen-retrieved using citrate buffer (Sigma Aldrich) for 25 minutes at 100°C. Slides were re-equilibrated to RT then washed in PBS for 10 minutes. If needed, Avidin/Biotin blocking kit (Vector, Newark, CA) was used as directed. Sections were blocked with Power Block (Biogenex, Fremont, California, USA) for 15 minutes at room temperature and washed twice in PBS. Slides were incubated in primary antibodies at 4°C overnight in 10% donkey serum in 0.1% Triton in PBS (PBSX). The following day, slides were washed in 3x (PBSX). Secondary antibodies (in 10% Donkey serum) were added to the slides and incubated at RT for 30 minutes. Slides were washed in 3x in PBSX and counterstained with DAPI (1:5000) for 10 minutes prior to a PBS wash and mounting in Prolong Gold (Invitrogen). Antibody details are outlined in **Supplemental Table 2**.

#### RNA-ISH

RNA-*in situ hybridization* was performed according to ACD (Advanced Cell Diagnostics) published protocols with the following probes: *Scgb1a1* (Cat No. 420351), *Scgb3a2* (Cat No. 809381), *Muc5b* (Cat No. 471991). For subsequent IF steps, after the final step of the protocol, slides were blocked with PowerBlock as above and additional immunofluorescent staining performed as above before nuclear staining with DAPI, cover slipping, and imaging.

#### Microscopy and image analysis

For immunofluorescence analysis, samples were imaged in a Keyence BZ-X710 inverted fluorescence microscope, images were taken on a 20X or 40x objective. Image analysis was performed using Halo (Indica Labs) Histological analysis of lung injury was quantified using ImageJ on samples stained with Masson’s Trichrome stain.

#### Mouse organoids

dTom+ cells were isolated via FACS as above and plated at 3000 cells/50 μL domed matrigel droplet in technical triplicate. Serum-free-feeder-free (SFFF) media conditions were used as described without modification. For organoids transitioned to Alveolar Differentiation media (ADM), those were cultured and expanded under SFFF conditions then switched to ADM media. Treatment with PT-2385 occurred upon plating until completion of the experiment at indicated concentrations.

#### Human organoids

Following CD326^+^ column-based enrichment, cells were plated into 50µl 1:1 v/v Matrigel droplets (Corning, Product Number 356231, Growth Factor Reduced (GFR) Basement Membrane Matrix, Phenol Red-free, LDEV-free) under Serum Free Feeder free conditions at a density of 25,000 cells/droplet (500 cell/μL) in either the Human SFFF media described previously by Katsura et al.(Katsura et al. 2020) (prepared within 2 weeks of use and stored at 4C), or in Alveolar Organoid Expansion Media (Stemcell Technologies Catalog #100-0847; according to manufacturer instructions including Antimycotic/Antibacterial). 10 µM ROCK-inhibitor Y-27632 (Tocris, catalog no. 1254) was added to initial plating media for the first 72 hours. If human alveolar differentiation media(ADM) was used, it was freshly prepared and used within 2 weeks per previously published protocol(*50*). Media was changed every 48-72 hrs for an initial outgrowth period of 2-3 weeks to enable robust expansion Passaging was performed either in a bulk fashion through removal of the domed droplet from the with a P1000 pipette, transfer to precooled 1.5 ml tube, addition of appropriate volume of splitting media, trituration with a p200 (unfiltered) over a P1000 with aggressive disruption, then addition of freshly thawed Martigel in a 1:1 v/v ratio, then replated. Alternatively, if single cell suspensions (for scRNA-seq) or passage were desired, the protocol from Jacob et al. (*90*) was used with minor changes where briefly, droplets containing organoids were digested using 2 mg/ml Dispase (dissolved in phenol-free DMEM with 25mM HEPES) for 1hr at 37C with interim disruption of matrigel after 30min with a P200 pipette. Ensuing suspension was washed 2x with ice cold DMEM with 25mM HEPES (300g for 5 minutes for intervening centrifugations). Following the second wash, all supernatant was manually aspirated (important to minimize residual Ca from media) and 500 μL/droplet (if done in bulk) of 0.05% Trypsin+EDTA was added and incubated at 37C for 7-12 minutes with an interval gentle triteration with a P1000 to aid in dissociation. Once a single cell suspension was obtained, an equal volume of ACF Enzyme Inhibition Solution (Stemcell Technologies, Cat. No. 05428) was applied and mixed, cells pelleted, washed 1x with 10 ml DMEM+ 25mM HEPES, and then used in downstream experiments.

#### Lysotracker Staining

Organoids were stained *in situ* and imaged in droplets. Lysotracker DeepRed (Invitrogen, Cat no. L12492) 75nM was applied in Optimem (Gibco Cat no. 31985062) for 1hr at 37C before returning to standard SFFFM.

### Data and code availability

Raw genomic data are available through the Gene Expression Omnibus (GEO) accession number (GSE243252). Code for genomic analysis is available at github.com/KropskiLab/Hif_2023.

### Statistical Analysis

Analysis performed in this work was analyzed using a combination of GraphPad Prism v9.5.1 and R v4.3.0. All statistical tests between groups were analyzed using non-parametric methods and specified in each figure. Specific analysis parameters associated with scRNA-seq data are outlined specifically within sections above along with rationale for their implementation and statistical diagnostics where applicable. Except where noted for clarity, all data are plotted as median ± 95% confidence intervals. For gene-related tests, multiple-testing adjusted p values were reported and used for thresholding.

## Acknowledgements

This work was supported by NIH/NHLBI R01HL145372 (JAK/NEB), R01HL153246(JAK), W81XWH1910415 (JAK/NEB), P01HL092870 (TSB), T32HL094296 (ASM), the Vanderbilt Faculty Research Scholars (JJG), and the Francis Family Foundation (JJG).

---

**Supplemental Table 1.**

| **Sample** | **Age** | **Sex** | **Diagnosis** | **Smoking**<br>**Ever** | **Current**<br>**(Pack yrs**<br>**total)** |
| --- | --- | --- | --- | --- | --- |
| HD4200 | 55 | M | Declined<br>Donor | Yes | No (5) |
| ILD146 | 60 | F | CTD-ILD | Yes | No (15) |
| ILD150 | 32 | M | uILD | Yes |  |
| ILD162 | 70 | F | IPF | Yes |  |

**Supplemental Table 2 –.**
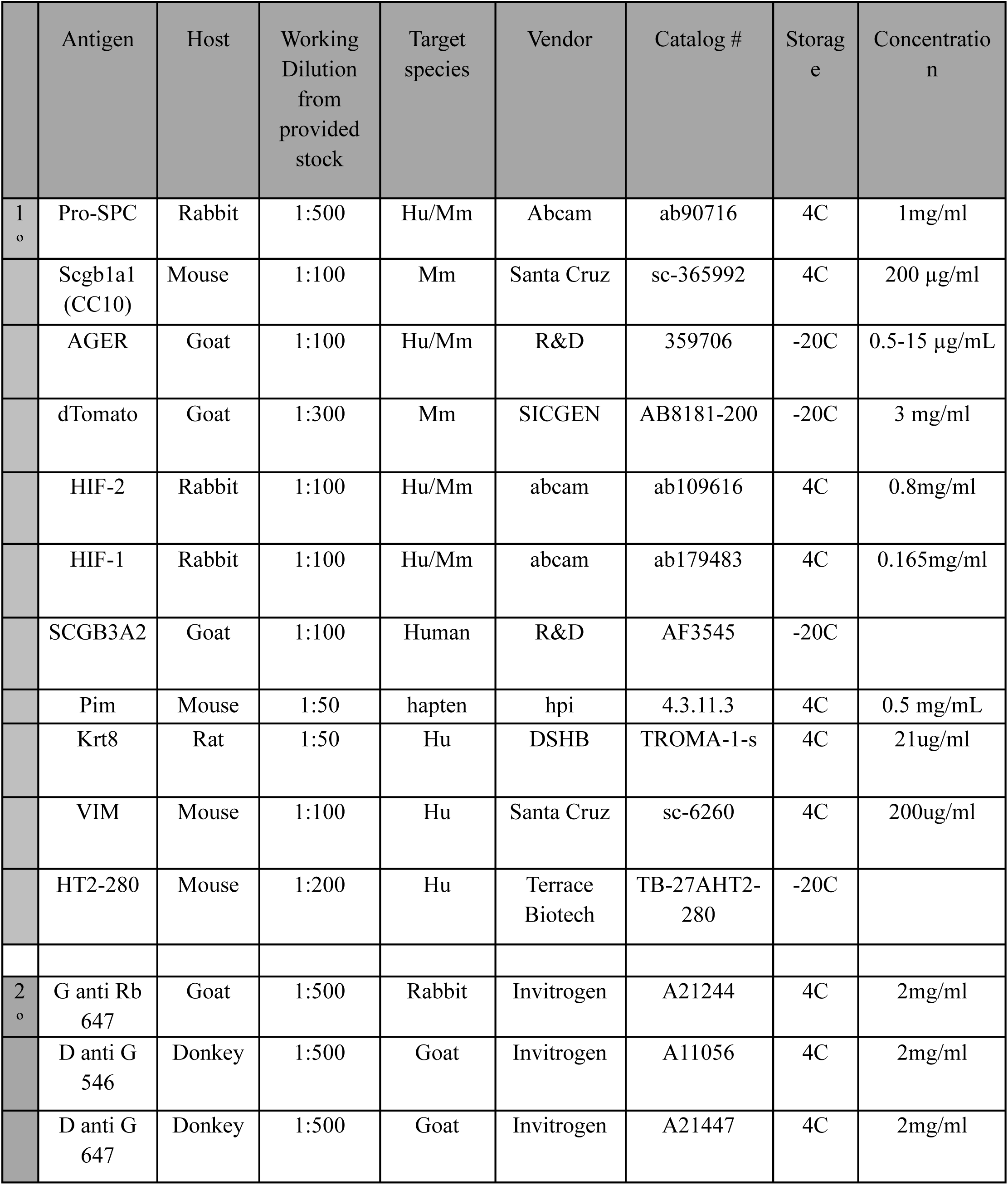
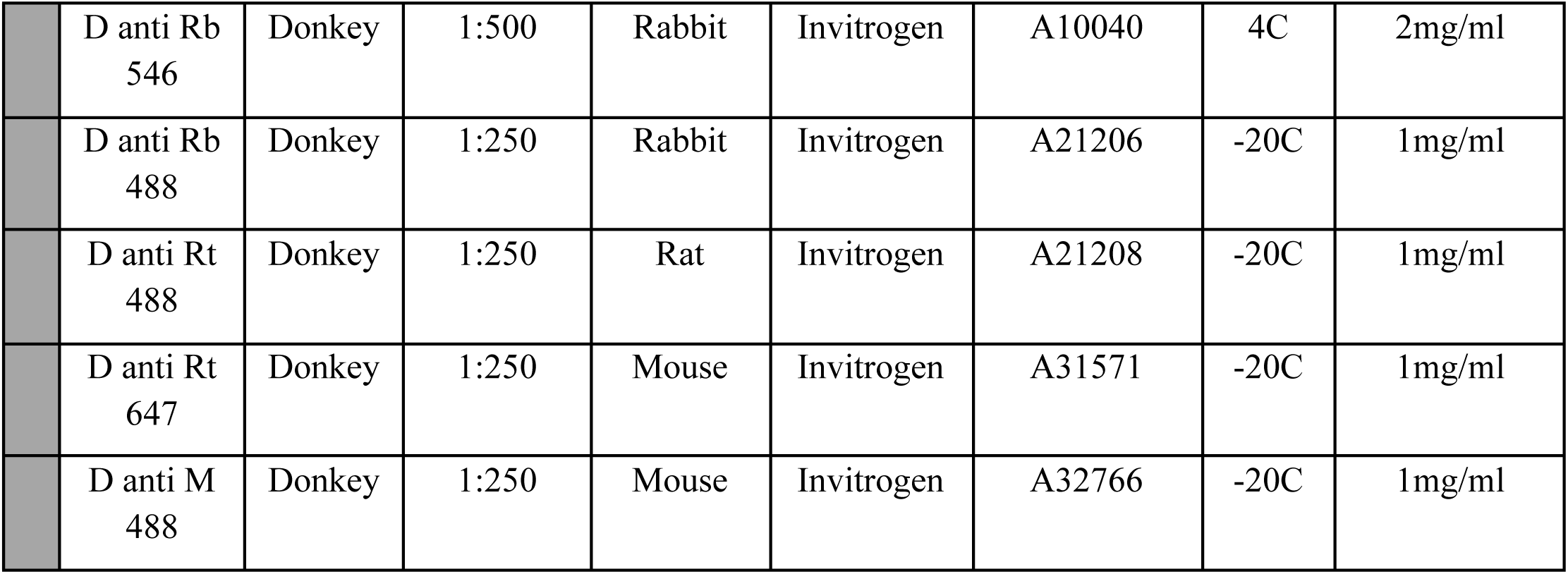
Antibodies used in Immunofluorescence imaging.

## Supplemental Figures

**Supplemental Figure 1-.**
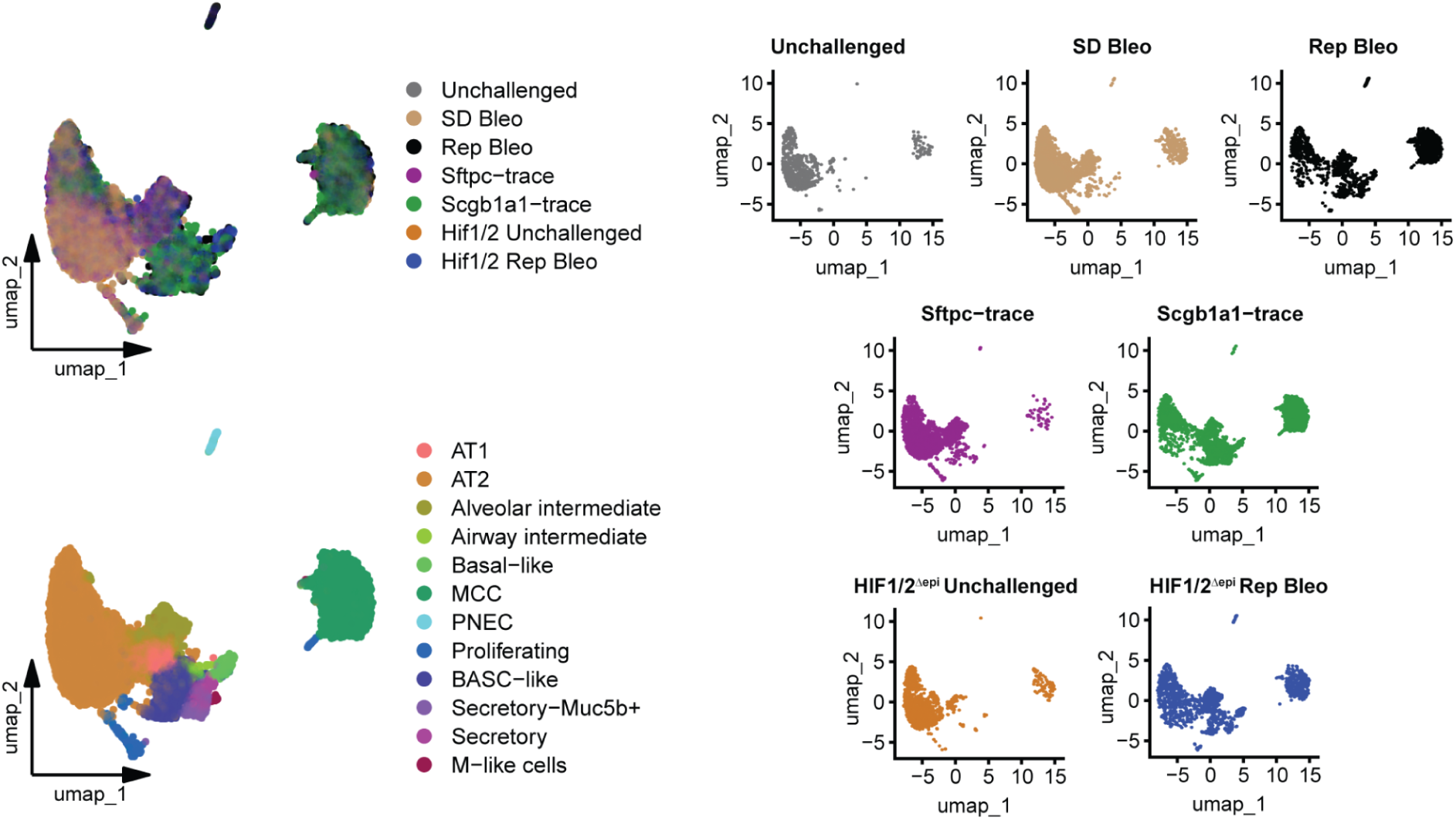
Joint embedding and annotation of murine scRNA-seq. Joint embedding and cell-type annotation across all murine data sets presented in Figures 1-3.

**Supplemental Figure 2-.**
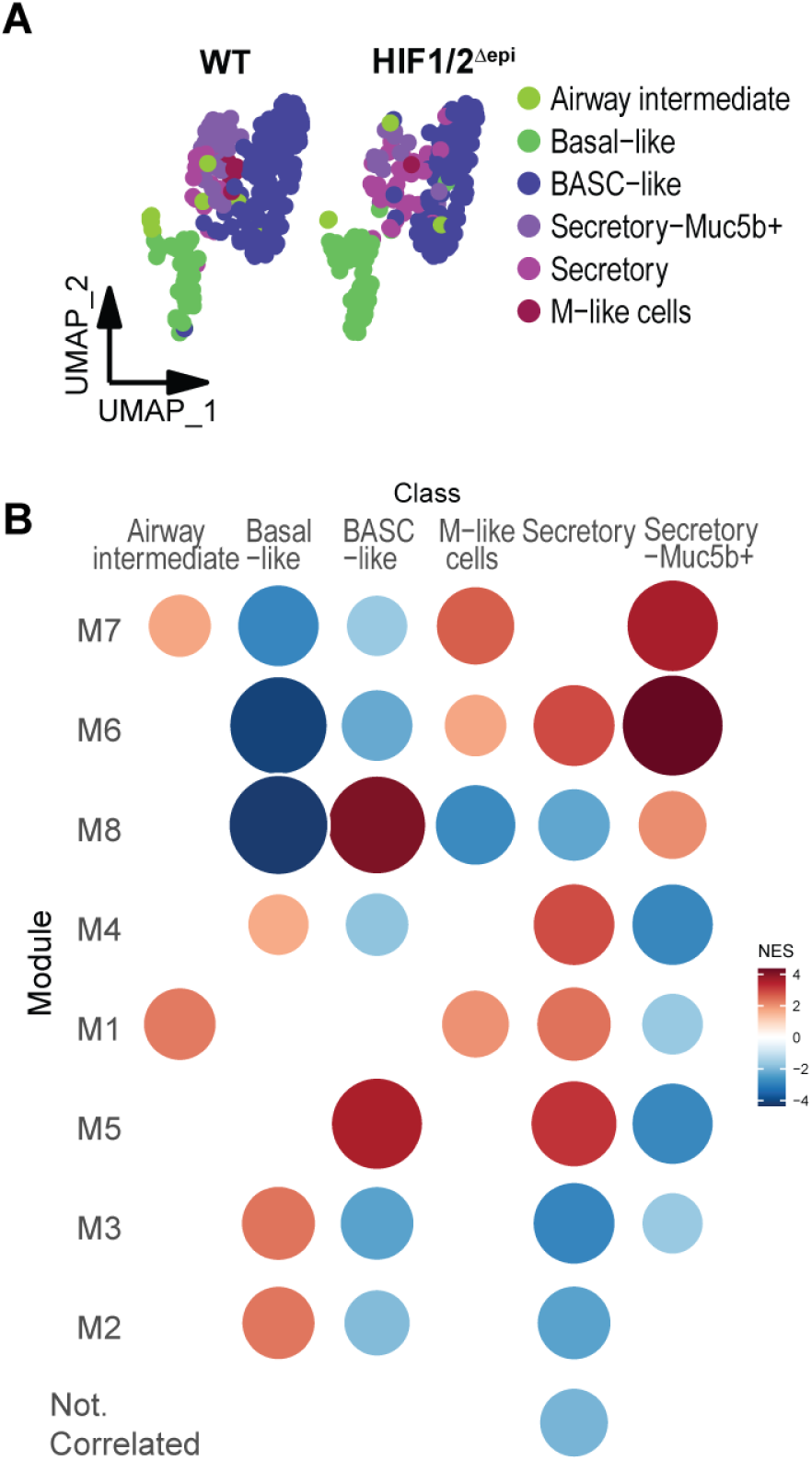
Secretory and airway basal cells analyzed as part of gene module identification. A) UMAP embedding of airway secretory cell and basal cell subset used in gene module analysis of Figure 2L. B) GSEA of same gene modules in Fig 2L based on scRNA-seq cell-type annotation plotted by Z score

**Supplemental Figure 3-.**
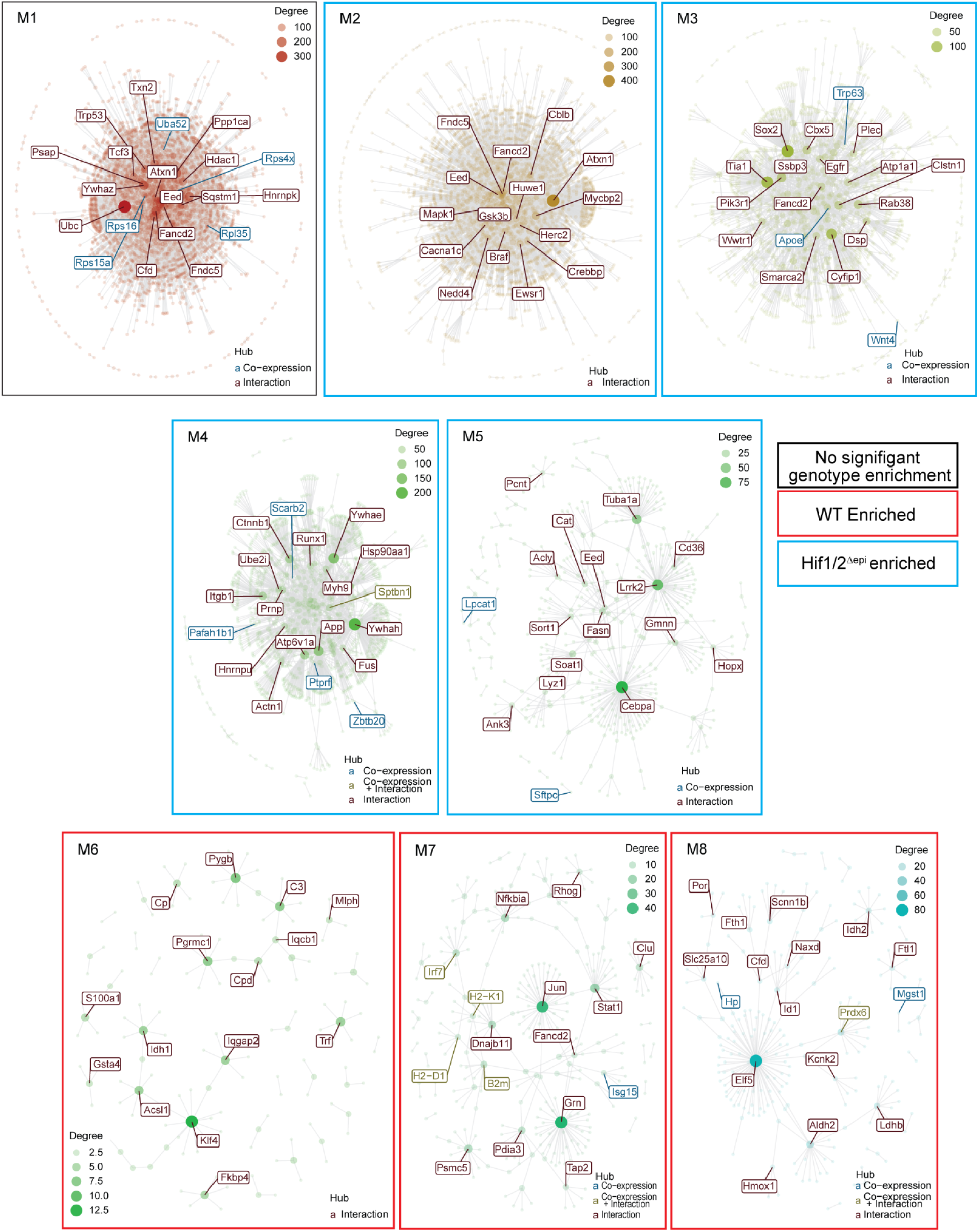
Protein-Protein interactions from airway gene modules. Gene modules (M1-M8) protein-protein interactions derived from the HitPredict(*82*) murine validated database.

**Supplemental Figure 4-.**
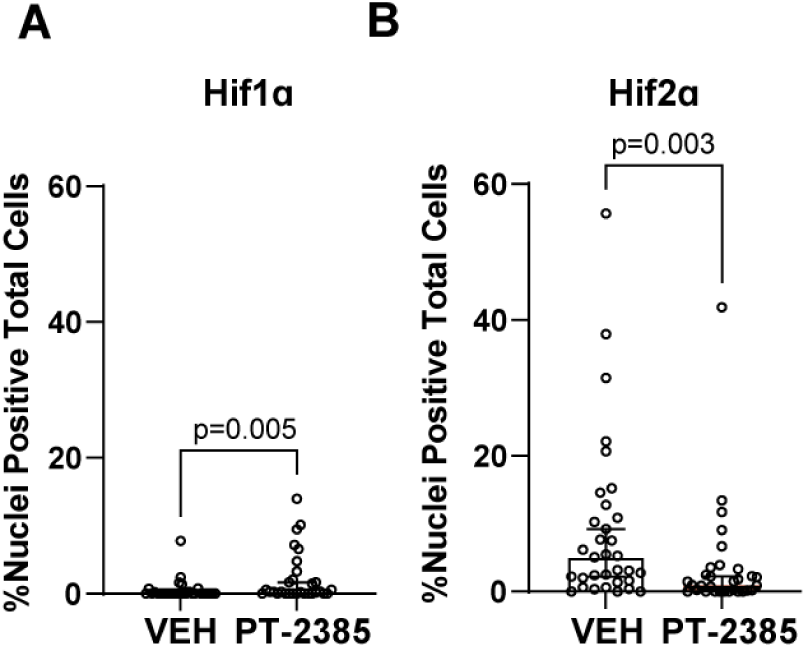
Nuclear Hif localization in Rep Bleo and inhibition of Hif2 nuclear localization with PT-2385 administration. A-B) Quantitation of Hif1α and Hif2α nuclear staining in dTomato^+^ cells in Vehicle and PT-2385 treated mice after Rep Bleo. Nuclear localization determined through automated image analysis (Halo, Indica Labs). N= 5 with 5 sections per sample. Plotted as median ± 95% CI

**Supplemental Figure 5 –.**
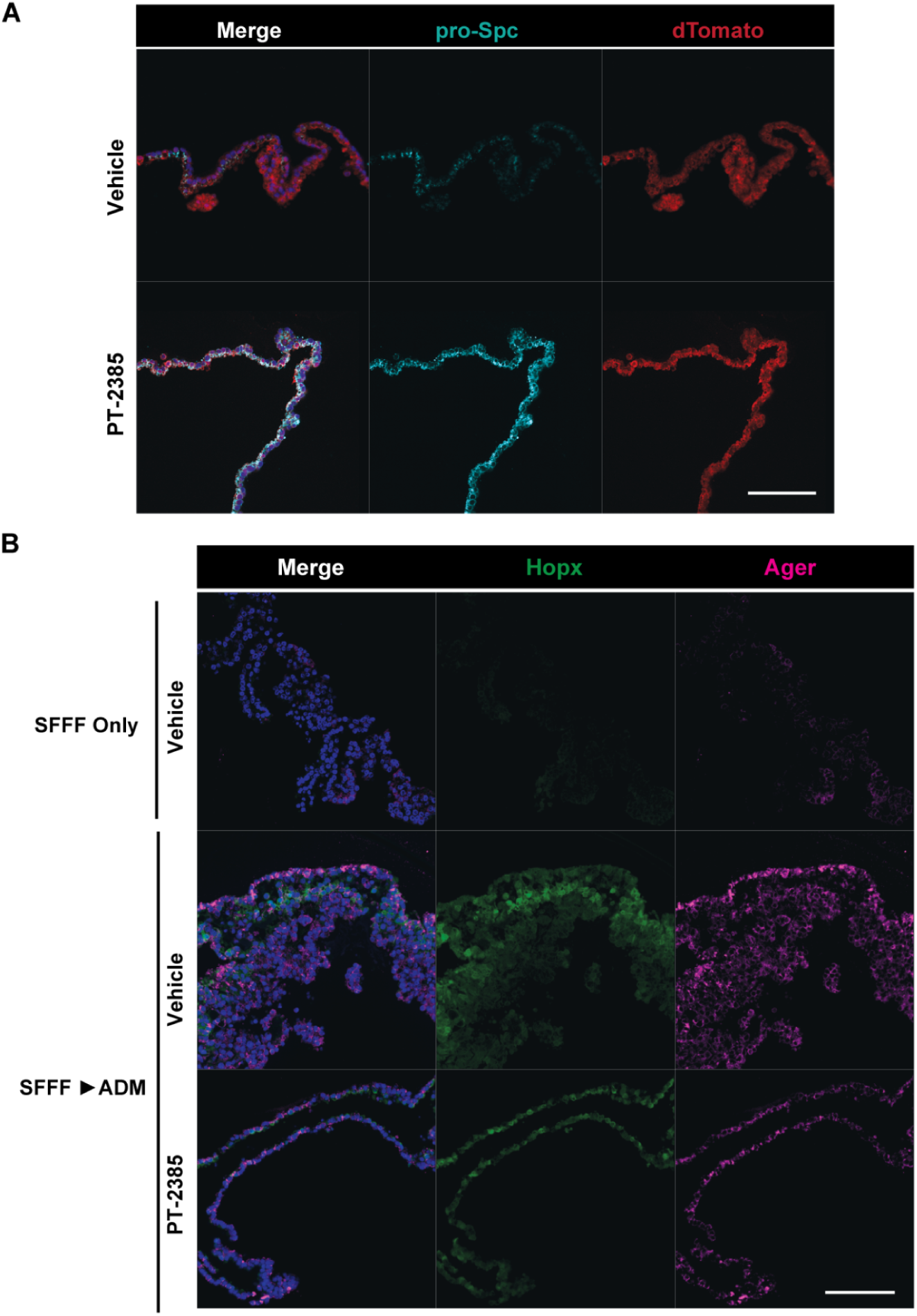
Expanded single channel immunofluorescence for airway derived alveolar organoids. A) Individual immunofluorescent channels corresponding to Figure 5C. Scale bar = 100 µm. B) Individual immunofluorescent channels corresponding to Figure 4E. Scale bar = 100 µm.

**Supplemental Figure 6.**
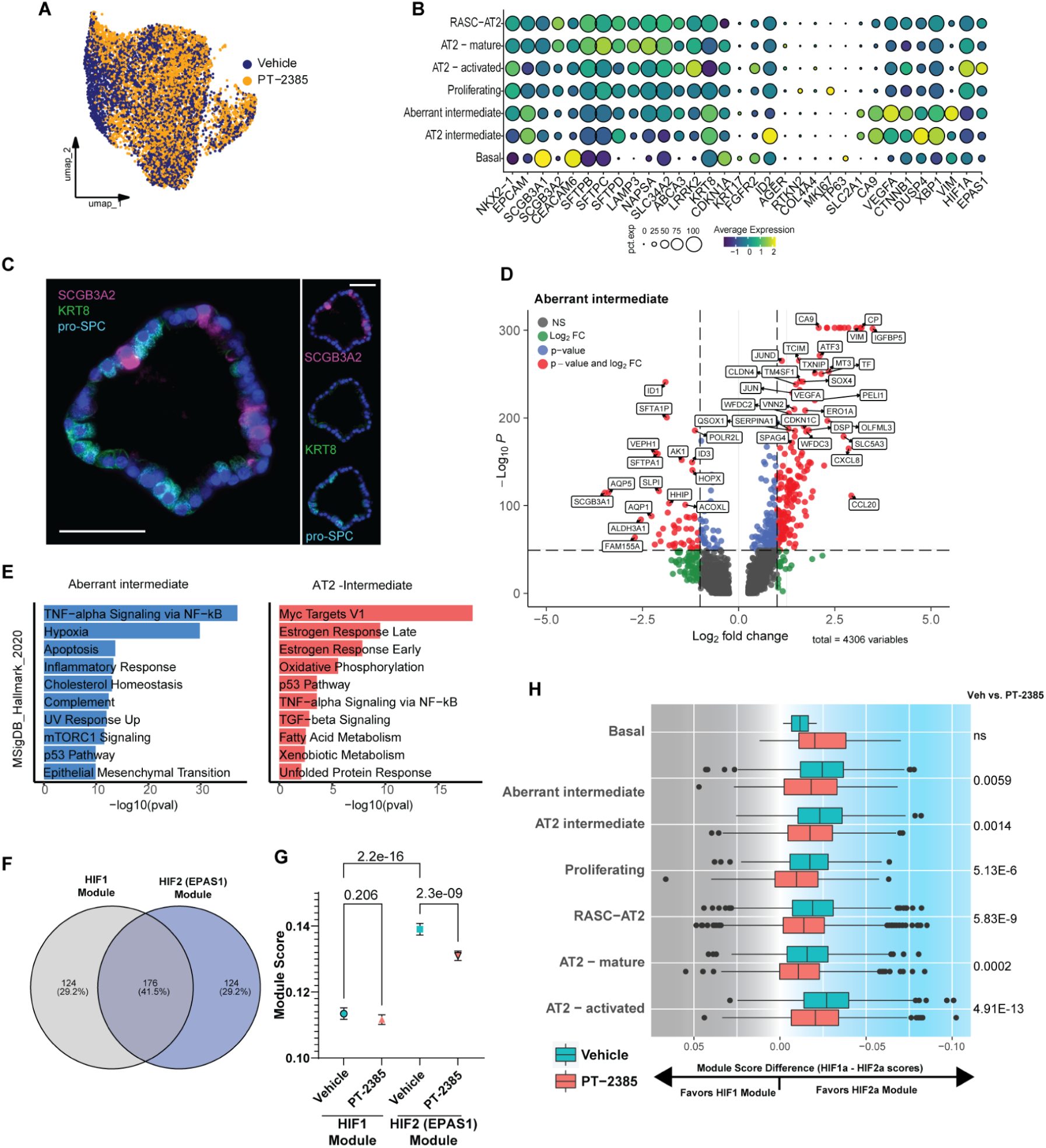
Human alveolar organoids exhibit activated hypoxia signaling. A) Dotplot of key markers for cell annotation. B) Representative IF image of SCGB3A2, pro-SPC, and KRT8 supporting the broadly observed scRNA proportions presented in Figure 6G. Scale Bar = 50 µm. C) Volcano plot for Aberrant intermediate differentially expressed (DE) genes. Plotted using EnhancedVolcano. D) Pathway analysis (MsigDB) comparing Aberrant Intermediate with AT2-intermediate clusters. E) Venn diagram showing overlap between validated HIF1 and HIF2 ChIP-seq targets across publically available data, obtained from Bono and Hirota(*83*). F) Plot of point estimates of HIF-specific module score by drug treatment demonstrated significant higher HIF2-targeted gene scores which were significantly decreased by PT-2385 treatment. Plotted as median ±95% CI. G) Bar plot comparing difference in HIF-module score (HIF1Score-HIF2Score) to assess individual cluster HIF-signal strength and effect from PT-2385 treatment. Pairwise p-values calculated and adjusted for multiple comparisons between Vehicle and PT-2385 for each cluster. PT-2385 treatment decreased HIF2-specific modules across each cluster.

**Supplemental Figure 7-.**
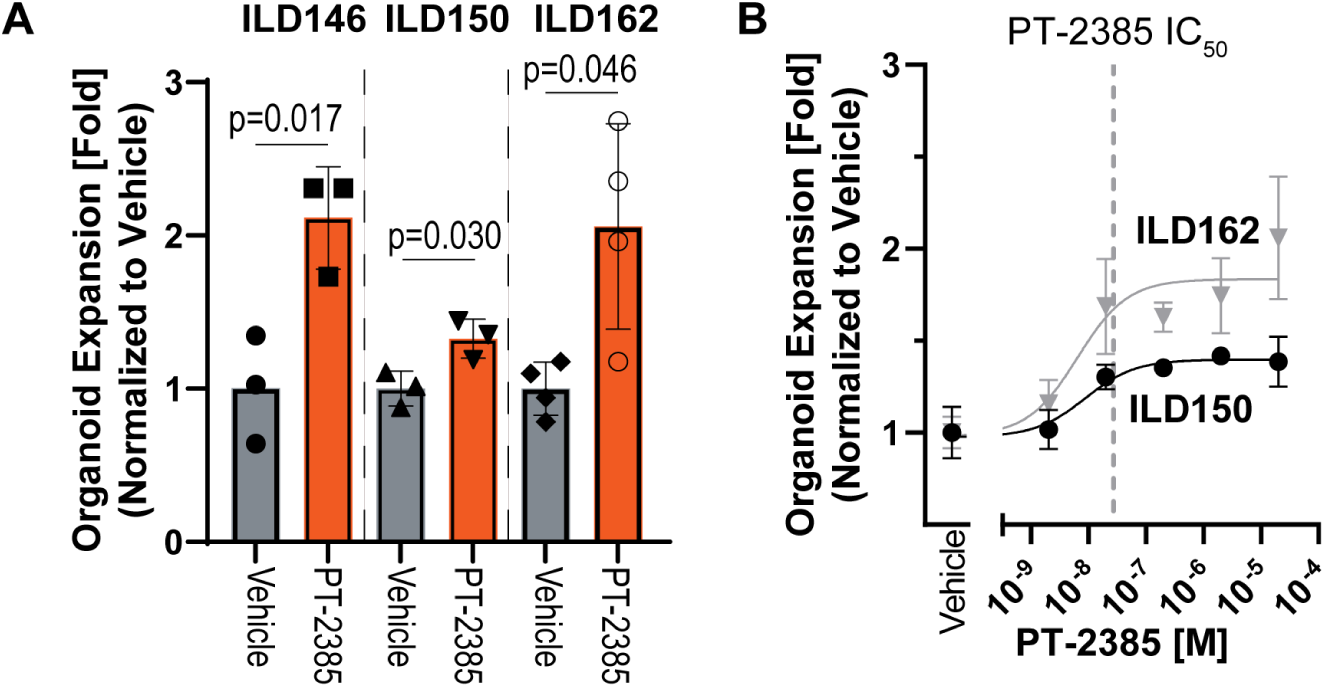
Dose-response of HIF2 inhibition on distal lung organoids from IPF-explanted lungs. A) Normalized organoid expansion of ILD-patient-derived explants following an initial 3-week outgroth and passage in SFFFM. B) Fitted dose-response for organoid growth on passage 1 organoids after 3-week initial outgrowth in SFFFM. Data plotted as mean ±SEM. EC50s found at 8.6 nM for ILD150 and 6.4nM for ILD162. Dotted line denotes published *in vitro* IC50 for PT-2385 on HIF2α:HIF1β dimerization. (*48*)

**Supplemental Figure 8-.**
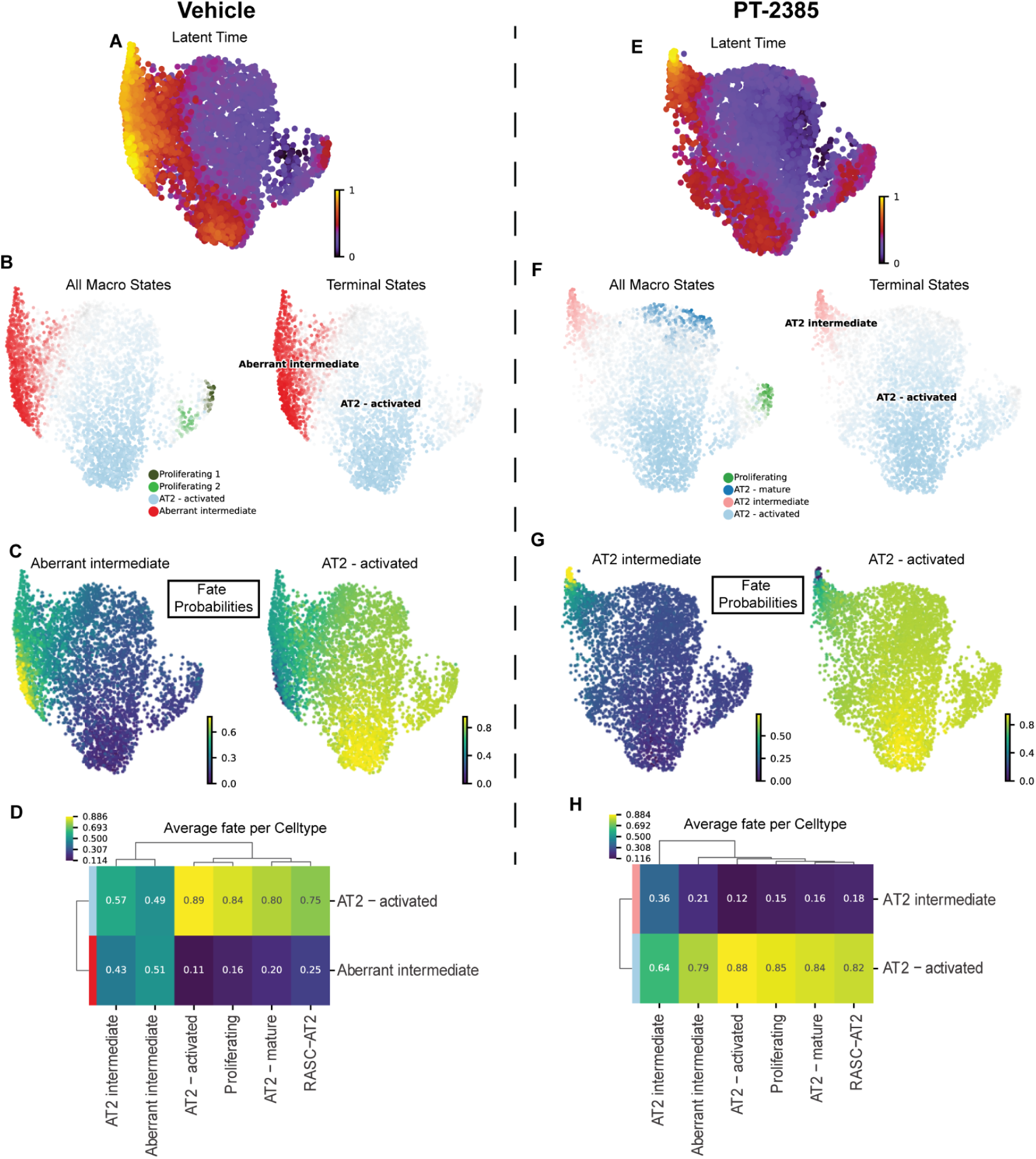
Differential RNA velocities and terminal states with HIF2-inhibition. Panels A-D represent Vehicle RNA velocity and E-H are separate PT-2385 treated organoid velocity analyzed separately to assess differential outcomes. A,E)UMAP embedding of Latent time calculation. B, F) Total macrostates and terminal states predicted using CellRank GPCCA estimator with default parameters and n=4 states based on Schur analysis of eigenvalues. C,G) UMAP of Absorption probabilities for predicted terminal states. D,H)Heatmap of absorption (Fate) probabilities based on cell cluster annotation

**Supplemental Figure 9-.**
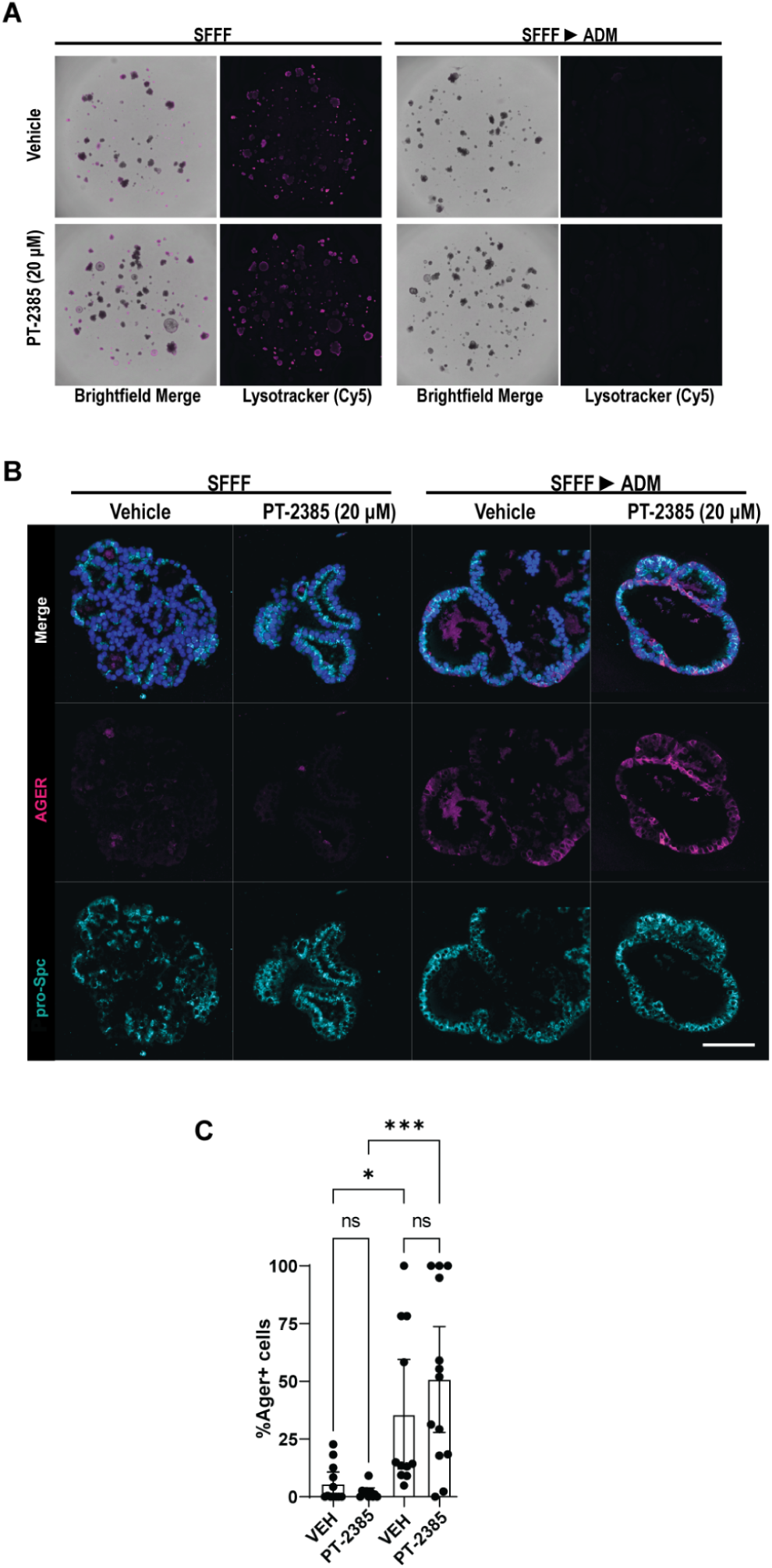
HIF2α inhibition does not adversely affect AT1 marker acquisition upon attempted differentiation. Representative images of Identical set of passage 2 organoids cultured in either SFFF only or transitioned to ADM after 10 days of culture and switched to ADM for additional 6 days. A) Brightfield and Lysotracker staining of human organoids in SFFF or transitioned to ADM demonstrating loss or lysotracker staining which is unaffected by PT-2385.B) Individual immunofluorescent channels. Scale Bar = 100 µm. C) Quantitation of IF of AGER and pro-SPC-positive cells following vehicle or PT-2385 treatment.

## Notes

### Competing Interest Statement

JAK reports grant funding from Boehringer Ingelheim, Bristol-Myers-Squibb. TSB reports grant funding from Boehringer Ingelheim, Bristol-Myers-Squibb and Morphic. NEB reports consulting fees from Deepcell.

